# Near-Infrared Afterglow Luminescence Amplification via Albumin Complexation of Semiconducting Polymer Nanoparticles for Surgical Navigation in Ex Vivo Porcine Models

**DOI:** 10.1101/2024.04.26.587487

**Authors:** Nathaniel Bendele, Ken Kitamura, Isabella Vasquez, Asma Harun, McKenzie Carroll, Indrajit Srivastava

## Abstract

Afterglow imaging, leveraging persistent luminescence following light cessation, has emerged as a promising modality for surgical interventions. However, the scarcity of efficient near-infrared (NIR) responsive afterglow materials, along with their inherently low brightness and lack of cyclic modulation in afterglow emission, has impeded their widespread adoption. Addressing these challenges requires a strategic repurposing of afterglow materials that improve on such limitations. Here, we have developed an afterglow probe, composed of bovine serum albumin (BSA) coated with an afterglow material, a semiconducting polymer dye (PFODBT/SP1), called BSA@SP1 demonstrating a substantial amplification of the afterglow luminescence (∼3-fold) compared to polymer-lipid coated PFODBT (DSPE-PEG@SP1) under same experimental conditions. This enhancement is believed to be attributed to the electron-rich matrix provided by BSA that immobilizes SP1 and enhances the generation of ^1^O_2_ radicals, which improves the afterglow luminescence brightness. Through molecule docking, physicochemical characterization, and optical assessments, we highlight BSA@SP1’s superior afterglow properties, cyclic afterglow behavior, long-term colloidal stability, and biocompatibility. Furthermore, we demonstrate superior tissue permeation profiling of afterglow signals of BSA@SP1’s compared to fluorescence signals using *ex vivo* tumor-mimicking phantoms and various porcine tissue types (skin, muscle, and fat). Expanding on this, to showcase BSA@SP1’s potential in image-guided surgeries, we implanted tumor-mimicking phantoms within porcine lungs and conducted direct comparisons between fluorescence and afterglow-guided interventions to illustrate the latter’s superiority. Overall, our study introduces a promising strategy for enhancing current afterglow materials through protein complexation, resulting in both ultrahigh signal-to-background ratios and cyclic afterglow signals.

## 1. Introduction

Currently, surgical procedures are the prevalent approach to treating various forms of solid tumors.^1^ The traditional methods to detect tumor margins rely upon palpitations and visual inspection, allowing surgeons to differentiate between healthy tissues and cancer tissue. However, using such information to guide surgical incisions has shown to be error-prone, mainly because many cancers aren’t palpable. Following surgery, this increases the risk of cancer recurrence.^2-4^ Hence, researchers have directed a lot of emphasis toward the development of preoperative and intraoperative imaging methods tailored toward specific characteristics of the tumor.^5,6^ This approach is crucial for helping surgeons remove as much tumor tissue as possible. In intraoperative imaging, fluorescence imaging stands out due to its high resolution, sensitivity, absence of radiation, and rapid feedback. In recent years, near-infrared (NIR) fluorescent dyes have emerged as promising tools for delineating tumor margins in both experimental cancer models and human patients.^7-9^ However, conventional probes in the NIR-I fluorescent window (having emission wavelengths between 700 and 900 nm) have limitations that include shallow tissue penetration (∼1–3 mm) and significant tissue autofluorescence, which diminish their effectiveness for fluorescence imaging-guided surgery (FIGS).^10,11^

Recently, afterglow luminescence, which refers to sustained luminescence after the cessation of excited light, has shown immense promise in advancing bioimaging applications ^12-16^, particularly in image-guided surgery and intraoperative imaging.^17-19^ Notably, afterglow imaging obviates the need for real-time excitation, effectively mitigating background signals stemming from tissue autofluorescence, a common limitation in NIR-I fluorescence imaging.^20,21^ Despite the advantages of afterglow luminescence for *in vivo* imaging, its widespread utility needs to be improved by the absence of NIR-responsive afterglow materials. The existing materials often exhibit suboptimal brightness and lack cyclic modulation, posing challenges to their broader adoption. Addressing these limitations necessitates a strategic reevaluation and repurposing of extant NIR-responsive afterglow materials to augment their efficacy in intraoperative surgical guidance.

Albumin, a significant protein in the body, plays a pivotal role in transporting diverse substances in the blood.^22^ Bovine serum albumin (BSA) closely resembles human serum albumin (HSA) with ∼80% sequence similarity, making it a cost-effective and readily purified carrier molecule. Its structural features enable it to bind to positively and negatively charged molecules, as well as hydrophilic and hydrophobic compounds.^23-28^ Notably, when fluorescent dyes, particularly cyanine dyes, are covalently conjugated or undergo supramolecular encapsulation with BSA, they have demonstrated several advantages, including (i) enhancing fluorescence brightness and photostability, (ii) improved colloidal stability by shielding against endogenous enzymatic degradation, and (iii) improved biocompatibility.^29,30^ Recent advancements have extended BSA’s utility to include coating aggregation-induced emission dyes,^31^ of which have been shown to improve their photodynamic therapy efficacy. This improvement arises from BSA providing an electron-rich environment that promotes the generation of the type-I reactive oxygen species. However, to date, no studies have explored the potential of a BSA coating to boost the afterglow luminescence of afterglow substrates.

Here, we have developed an afterglow probe, BSA@SP1, which consists of bovine serum albumin (BSA) coated with a semiconducting polymer dye, Poly[2,7-(9,9-dioctylfluorene)-*alt*-4,7-bis(thiophen-2-yl)benzo-2,1,3-thiadiazole] (PFODBT/SP1). Specifically, we show that BSA coating imparts BSA@SPs with superior afterglow luminescence properties when compared to conventional polymer encapsulation matrix, 1,2-Distearoyl-sn-glycero-3-phosphoethanolamine-Poly(ethylene glycol) (DSPE-PEG) by ∼3-fold, and cyclic afterglow behavior. We hypothesize such an enhancement could be attributed to the electron-rich matrix provided by BSA, which immobilizes PFODBT and enhances its generation of ^1^O_2_ radical generation, thus improving the afterglow luminescence brightness. Our preliminary molecular docking studies reveal that PFODBT preferentially inserts into the hydrophobic pocket of BSA, imparting BSA@SP1 nanoparticles with improved colloidal stability and biocompatibility. Moreover, we showcase enhanced tissue permeation profiling by leveraging the afterglow capabilities of BSA@SP1 through experiments conducted on *ex vivo* tumor cell-mimicking phantoms^32,33^ and various porcine tissue types (including skin, muscle, and fat), revealing their superior signal-to-background ratio compared to fluorescence imaging. Expanding on this, to assess its effectiveness in image-guided cancer surgeries, we implanted tumor-mimicking phantoms at various locations within the porcine lungs: (i) 0.5mm below the right inferior lobe, (ii) 5mm below the left inferior lobe, and (iii) within the thoracic cavity. Subsequently, we conducted direct comparative studies between fluorescence- and afterglow-guided surgical interventions, highlighting the superior performance of afterglow-assisted procedures. In summary, our approach of protein immobilization to enhance afterglow luminescence in existing materials holds significant promise, as it improves signal-to-background ratios, benefits from a low afterglow background, and exhibits cyclic afterglow luminescence. These attributes showcase how BSA modification of afterglow materials enhances their imaging capabilities and serves as a highly promising candidate for image-guided cancer surgery in *in vivo* models.

## 2. Results and Discussion

### 2.1. Synthesis of BSA@SPs

For the development of BSA@SP1 nanoparticles, a modified nanoprecipitation method was used (Figure 2a). SP1 (or PFODBT) was dissolved in THF and quickly added dropwise to BSA containing PBS1x solution under continuous probe sonication. For the development of our control nanoparticle, i.e. amphiphilic polymer (DSPE-PEG2k) coated SP1, i.e. DSPE-PEG@SP1, a general nanoprecipitation method was used. Both DSPE-PEG2k and PFODBT were co-dissolved in THF and quickly added to PBS1x solution under continuous probe sonication. Once the post-processing that involved centrifugation wash steps was completed, it led to clear nanoparticle solutions (Figure 2b, c: insets) for use in further physicochemical and optical characterizations.

**Figure 1.**
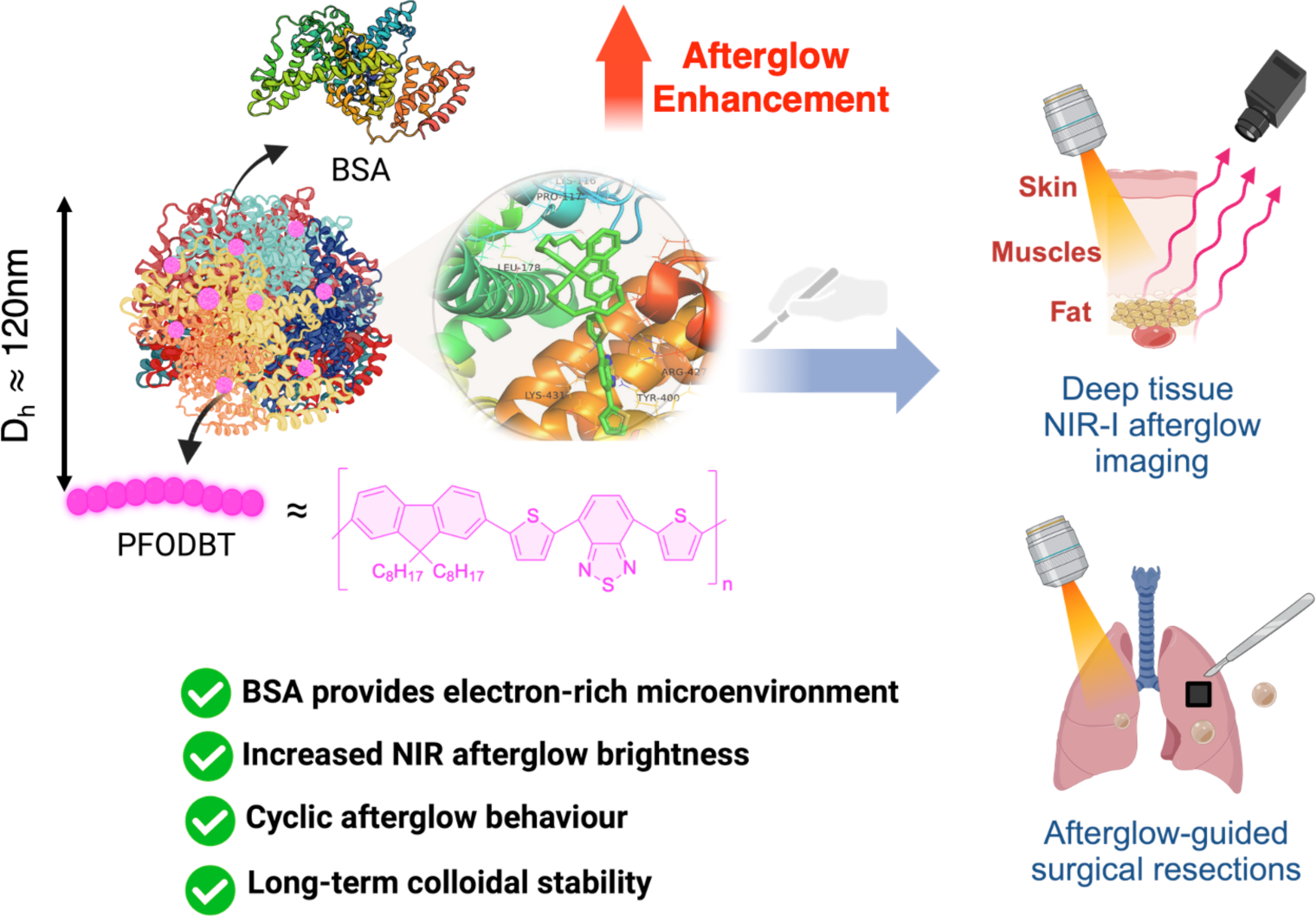
The schematic diagram illustrates the application of bovine serum albumin-coated PFODBT/SP1 semiconducting polymer nanoparticles (BSA@SP1). BSA@SP1 exhibits enhanced afterglow brightness and retention, long-term photostability, and colloidal stability. BSA@SP1 is further leveraged for deep tissue imaging and afterglow-guided surgical resection.

**Figure 2.**
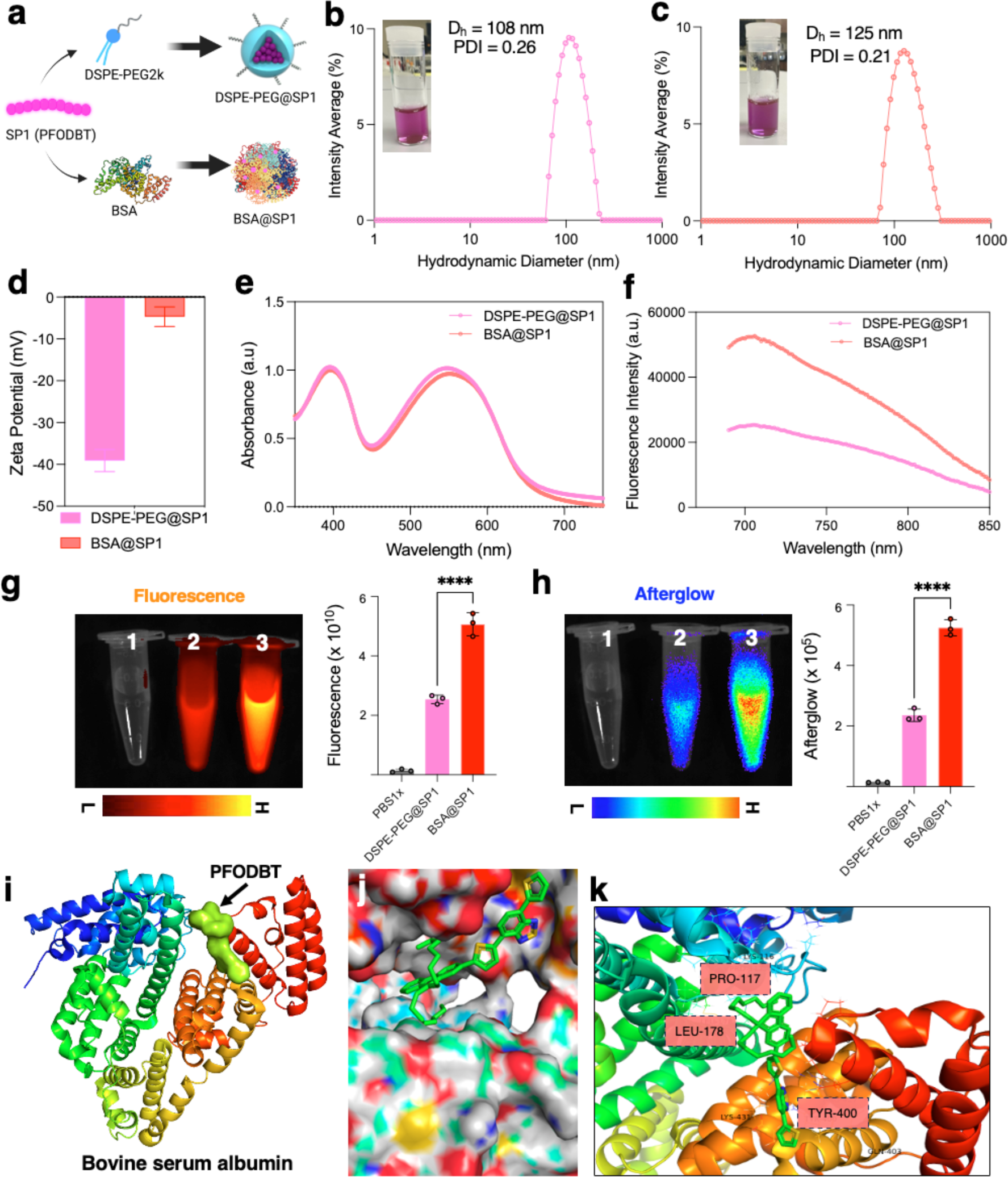
Physico-chemical and photophysical characterization of DSPE-PEG@SP1 and BSA@SP1. (a) Synthesis scheme of preparation of DSPE-PEG@SP1 and BSA@SP1 nanoparticles. Corresponding hydrodynamic diameter measurements from dynamic light scattering and insets showing the nanoparticle solutions for (b) DSPE-PEG@SP1, and (c) BSA@SP1, respectively. Zeta potential (d), UV-Vis-NIR absorption spectra (e), and fluorescence emission spectra at 645nm excitation (f) of DSPE-PEG@SP1 and BSA@SP1. Fluorescence images (g) and afterglow images (h) were acquired using an IVIS imager and their corresponding intensity values were calculated with 1: PBS1x, 2: DSPE-PEG@SP1, 3: BSA@SP1. Molecular docking studies to evaluate the binding between PFODBT and BSA and analyzed with : (i) Receptor (BSA) as a cartoon and ligand (PFODBT) as a stick. (j) Receptor (BSA) as a surface and ligand (PFODBT) as a stick. (k) Interaction of the docked ligand (PFODBT) with the surrounding hydrophobic amino acid residues in BSA (highlighted in red).

### 2.2. Physicochemical and Optical Characterization of BSA@SPs

The hydrodynamic diameter of DSPE-PEG@SP1 and BSA@SP1 were measured using dynamic light scattering (DLS) and showed sizes of ∼108nm and ∼125nm, respectively (Figure 2b,c). They both further had low polydispersity index (PDI) values which indicated a narrow size distribution and particles being relatively uniform in size. Transmission electron microscopy (TEM) images were acquired to reaffirm their spherical morphology (Figure S1). Subsequently, ζ-potential measurements were performed, showing ∼ -38mV for DSPE-PEG@SP1. However, this decreased drastically for BSA@SP1 to ∼ -6mV (Figure 2d, Figure S2). A plausible explanation could be attributed to the electrostatic screening effect by BSA molecules, effectively screening the original surface charge of the nanoparticles. Hydrodynamic diameter and zeta potential values were obtained for both DSPE-PEG@SP1 and BSA@SP1 for 1 week while stored at 4°C (Figure S3). Long-term colloidal stability of BSA@SP1 was evaluated at (i) 37°C (to mimic physiological temperature) and (ii) 10% plasma solution (to mimic biological milieu). They demonstrated no significant change in the hydrodynamic diameter or PDI values, indicating their size stability and homogeneity (Figure S4). During this time, no visible precipitations were observed, indicating their excellent colloidal stability. Next, optical characterizations were conducted for both DSPE-PEG@SP1 and BSA@SP1 dispersed in PBS1x buffer, with a pH of 7.4. The absorption spectra revealed a prominent absorbance peak at 535nm for both nanoparticles, consistent with previous reports and our bandgap measurements.^32^ Upon excited at 605nm, both nanoparticles exhibited an emission peak at 820nm. However, BSA@SP1 displayed a significantly higher emission intensity compared to DSPE-PEG@SP1. To further investigate their fluorescence properties, we captured fluorescent images using the In Vivo Imaging System (IVIS) imager with excitation at 605nm and varying emission wavelength (680nm – 840nm). The fluorescence intensities demonstrated a similar trend for both nanoparticles (Figure 2g and Figure S5). Additionally, we acquired afterglow images using the bioluminescence mode of the IVIS imager after exiciting it with white light irradiation. Remarkably, BSA@SP1 showed substantially higher afterglow luminescence compared to DSPE-PEG@SP1 (Figure 2h). Both fluorescence and afterglow images of SP1/PFODBT (i.e. precursor molecule) were acquired using the IVIS Imager for reference (Figure S6). The hydrophobic nature of PFODBT allows it to interact with BSA and become embedded within its hydrophobic domains. To investigate this interaction and its underlying effect on the improved optical behavior of PFODBT in the presence of BSA, preliminary molecular docking studies were conducted between PFODBT and BSA. As shown in Figure 2i-k, the amino acid residues of BSA exhibited strong hydrophobic interaction with PFODBT, particularly hydrophobic amino acids such as PRO-117 and LEU-178, which interacted with the alkyl side chains of PFODBT. Moreover, the aromatic ring of TYR-400 formed π − π stacking interactions with the aromatic ring of PFODBT. These close interactions potentially limit the motion of PFODBT, thereby suppressing non-radiative decay and promoting the intersystem crossing process, ultimately enhancing the ^1^O_2_ radical generation efficacy of PFODBT in the presence of an electron-rich environment of BSA. These factors culminate into enhanced afterglow brightness for BSA-coated PFODBT nanoparticles, compared to traditional encapsulation matrices like DSPE-PEG, which do not have any effects on the optical properties of PFODBT.

### 2.3. Evaluation of Afterglow Emission Characteristics of BSA@SPs

Following an exhaustive physicochemical characterization and a thorough assessment of the long-term colloidal stability of BSA@SP1, our objective was to optimize the afterglow emission of BSA@SP1. We tested a range of concentrations for our BSA@SP1 nanoparticles, from 1 to 200 μg/mL. Our findings revealed a clear and strong correlation between the concentration of nanoparticles and the intensity of afterglow luminescence signals (Figure S7). We employed varying concentrations of BSA (0.66, 6.6 mg, and 13.2 mg/ mL) to coat the SP1 particles, forming BSA@SP1 nanoparticles. Subsequently, we quantified the afterglow emission intensities, which revealed the highest afterglow luminescence for the BSA@SP1 nanoparticles formed with 6.6 mg/ mL BSA (referred to as BSA66@SP1), while the emission from the BSA@SP1 nanoparticles formed with 13.2 mg/ mL BSA (referred to as BSA132@SP1) was comparable (Figure S8). Conversely, we observed the lowest emission for the complex formed with 0.66 mg/ mL BSA. Notably, as BSA132@SP1 exhibited a smaller dynamic light scattering (DLS) peak (∼10 nm), likely due to excess BSA that could not be removed from centrifugation wash steps it posed a potential risk of instability for future experiments (Figure S9). Consequently, to ensure stability and reliability in subsequent measurements, we opted to use BSA66@SP1, further referred to as simply BSA@SP1. For our control nanoparticle, DSPE-PEG2k was used as a surfactant to coat SP1 using nanoprecipitation.

Next we attempted to investigate how varying duration of light irradiation impacts the NIR afterglow luminescence emission of BSA@SP1. We observed that the intensity of the NIR afterglow from BSA@SP1 gradually increased with longer irradiation times, reaching its peak at 30 seconds and maintaining a similar signal for 60 seconds of irradiation (Figure 3b,e). In contrast, the fluorescence signal emitted by BSA@SP1 remained nearly constant regardless of the duration of irradiation (Figure 3f). This indicates that while the irradiation activates NIR afterglow luminescence for BSA@SP1, it does not affect their fluorescence emission. The irradiation had no impact on the hydrodynamic diameter or polydispersity index of BSA@SP1 (Figure S10). A similar trend was noted for DSPE-PEG@SP1, albeit with weaker afterglow luminescence and fluorescence emission signals compared to BSA@SP1 (Figure 3a,c,d).

**Figure 3.**
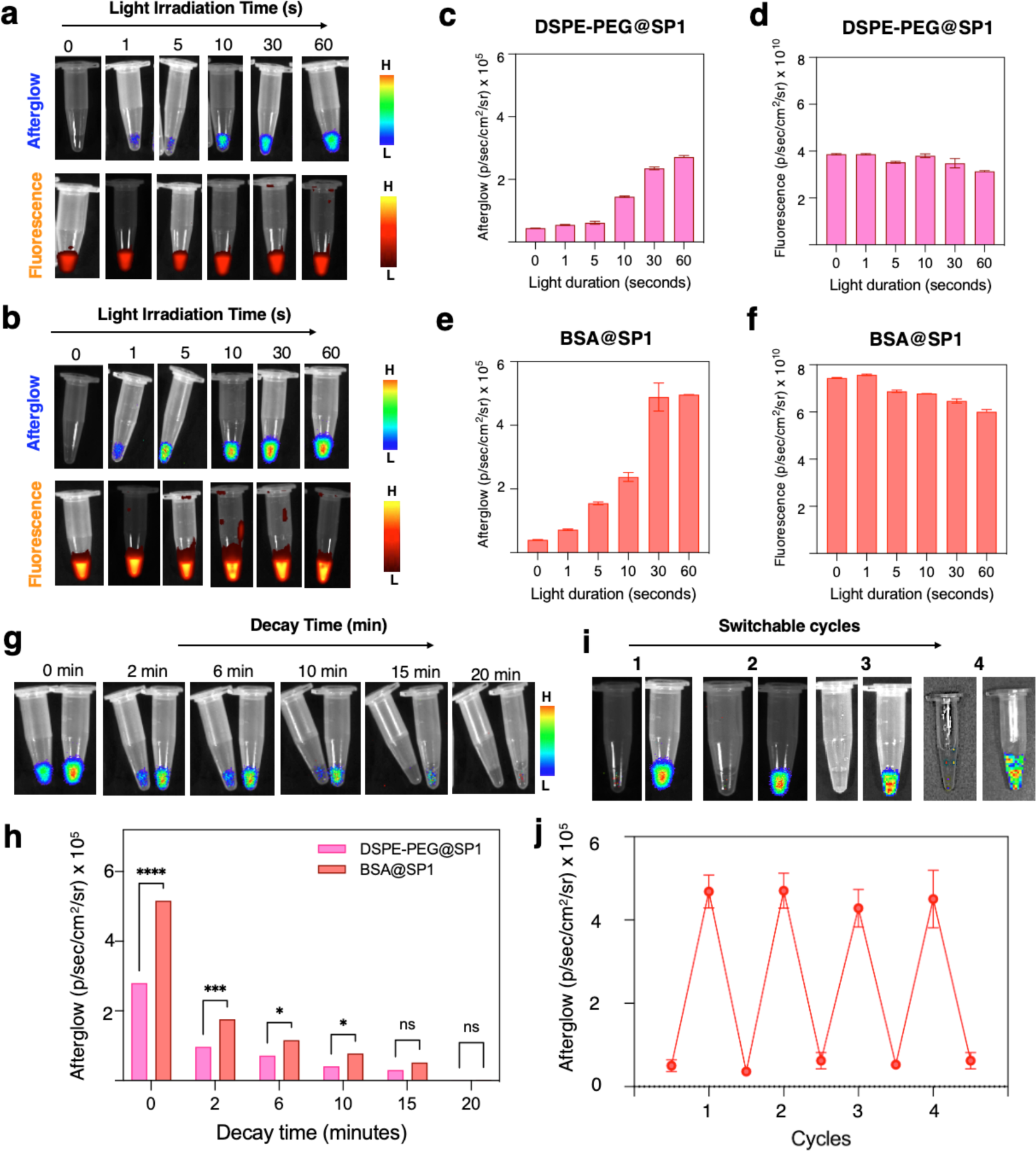
Evaluation of enhanced afterglow luminescence of BSA-coated PFODBT nanoparticles, i.e. BSA@SP1. Afterglow and fluorescence images of DSPE-PEG@SP1 (a) and BSA@SP1 (b) after receiving different light irradiation times and their corresponding afterglow (c, e) and fluorescent intensities (d, f) being calculated. Luminescence decay studies were conducted for DSPE-PEG@SP1 (left) and BSA@SP1 (right) and their afterglow images (g) and intensities (h) were calculated after receiving irradiation. (i, j) In each cycle, the nanoparticles were irradiated with light and then allowed to release photons for 20 minutes. The total duration of each cycle, which includes both the irradiation and the photon release periods, was 80 minutes of experimental time.

We aimed to investigate the persistence of NIR afterglow emission in BSA@SP1 once light irradiation ceased, as this feature could extend imaging time without compromising quality. Dynamic NIR afterglow images of BSA@SP1 revealed luminescence persisting for approximately 15 minutes post-irradiation (Figure 3g). In contrast, DSPE-PEG@SP1 exhibited weaker persistence, with luminescence fading within approximately 8 minutes. We conducted a lifetime analysis of the afterglow luminescence decay for DSPE-PEG@SP1 (t_1/2_ = 0.9 minutes) and BSA@SP1 (t_1/2_ ∼ 2 minutes), demonstrating that a BSA coating enhances the half-life (Figure S11). Further refinement could involve optimizing the ratio of SP to BSA to potentially extend this lifetime. Previous research has explored incorporating multi-components^47^ into afterglow nanoparticles to further enhance the afterglow lifetime, offering avenues for adaptation.

Subsequently, we attempted to investigate if BSA@SP1 could retain its afterglow luminescence across multiple cycles of irradiation. Such a characteristic would indicate their stability and durability, making them advantageous for prolonged observation for *in vivo* imaging without the need for repeated administration of the afterglow agent, thereby minimizing perturbations to the in vivo system under investigation. BSA@SP1 was still bright after four rounds of irradiation (Figure 3i) and 80 minutes of experimental time.

To elucidate the underlying mechanisms dictating the NIR afterglow luminescence in BSA@SP1, we first focused on whether heating could induce SP1/PFODBT to emit NIR afterglow. However, upon heating up to 60°C, we observed no discernible NIR afterglow luminescence. In contrast, exposure of BSA@SP1 to 60 seconds of white light irradiation resulted in a notable enhancement of NIR afterglow intensity (Figure S12). Subsequently, we subjected BSA@SP1 to both hypoxic and normoxic environments and captured their respective afterglow luminescence images. Remarkably, under hypoxic conditions, the NIR afterglow luminescence of BSA@SP1 was significantly diminished compared to that under normoxic conditions (Figure S13). To demonstrate that the BSA protein was indeed responsible for the afterglow enhancement, we denatured the BSA solution by heating it at 100°C for 20 minutes to diminish its functionality. We then used this denatured BSA to coat PFODBT/SP1 via nanoprecipitation. The resulting nanoparticles (dBSA@SP1) did not exhibit the same afterglow enhancement as BSA@SP1, where the BSA remains functional and provides an electron-rich environment (Figure S14). Furthermore, since most solid tumors are weakly acidic, we investigated the behavior of BSA@SP1 nanoparticles at acidic pH levels (∼4.5-5). When BSA@SP1 was incubated in PBS buffers at different pH values (4.5, 5, 5.5, and 6), the afterglow intensity was highest at pH 4.5 (Figure S15). This could be due to the increased generation of ^1^O_2_ radicals at lower pH levels.^34^

This observation implies that ^1^O_2_ radicals might play a role in modulating the NIR afterglow luminescence of BSA@SP1.^35^ Consequently, the logical progression was to investigate the generation of ^1^O_2_ radicals, as earlier studies have linked their production to the afterglow phenomenon in PFODBT or SP1.^35-38^ To assess the generation of ^1^O_2_ radicals from SP1, we used a singlet oxygen sensor green (SOSG). After the white light irradiation was increased (1s to 60s), the fluorescence intensity of SOSG at 525nm increased by ∼1.5x to 4x for BSA@SP1, as shown in Figure S16, suggesting the role the generation of ^1^O_2_ radical plays afterglow luminescence. Interestingly, when exposed to white light irradiation, the fluorescence intensity of SOSG at 525nm was significantly higher for BSA@SP1 than DSPE-PEG@SP1 (Figure S17).

A plausible explanation for this effect could be attributed to BSA’s role in providing a rich electron microenvironment, which acts as a substrate for ^1^O_2_ radical generation.^31^ Next, we investigated the correlation between NIR afterglow luminescence intensity and the generation of singlet oxygen in solution. As the irradiation time increased from 1s to 60s, we observed a substantial enhancement in the afterglow intensities of BSA@SP1. This was positively related to the increase in the fluorescence emission intensity of SOSG, indicating a corresponding rise in the amount of ^1^O_2_ radicals. Remarkably, we observed a strong correlation (Figure S18) between the yield of ^1^O_2_ radicals and NIR afterglow luminescence intensity (R^2^ = 0.92). To gain deeper insights into the molecular mechanisms underlying the heightened generation of singlet oxygen (^1^O_2_) observed in BSA@SP1 and its subsequent afterglow luminescence, we conducted comparative experiments utilizing another SP dye, PCPDTBT, designated as SP2 (Figure S19). By functionalizing SP2 with DSPE-PEG (DSPE-PEG@SP2) or BSA (BSA@SP2), we noted a comparable increase in ^1^O_2_ production using the SOSG assay, with a greater enhancement observed for BSA compared to DSPE (Figure S20). However, both DSPE-PEG@SP2 and BSA@SP2 exhibited fluorescence (Figure S21) but no discernible afterglow luminescence (Figure S22). This observation suggests a potential influence of the thiophene moieties present in SP1/PFODBT structures driving the afterglow luminescence, which is absent in SP2/PCPDTBT structures (Figure S19).

### 2.4. *In Vitro* Cytotoxicity Assessment and Cellular Internalization of BSA@SP1 in Cancer Cell Lines

After thoroughly characterizing BSA@SP1 and DSPE-PEG@SP1, we assessed their biocompatibility and ability to be internalized by 4T1 tumor cells *in vitro*. Both DSPE-PEG@SP1 and BSA@SP1 demonstrated minimal toxicity to the 4T1 cells, even at high concentrations of 100 µg/mL (Figure S23). To examine cellular internalization, we utilized flow cytometry to quantify uptake into 4T1 cells. Both DSPE-PEG@SP1 and BSA@SP1 exhibited increased uptake compared to the control (PBS 1x treatment). Particularly, BSA@SP1 demonstrated significantly higher cellular uptake (∼27%) compared to DSPE-PEG@SP1 (∼9.5%) (Figure S24). This trend may be attributed to the presence of secreted proteins acidic and rich in cysteine (SPARC) receptors that are highly expressed on metastatic tumor cells, such as 4T1, which are known to bind to albumin.^39^ These receptors likely facilitate a selective endocytosis uptake mechanism for BSA@SP1. To test our hypothesis, we first attempted to oversaturate the SPARC receptors in 4T1 cells by pre-treating them with albumin-rich media (100 µM BSA in PBS1x). Following this pre-treatment, the cells were incubated with 100 µg/mL of BSA@SP1 nanoparticles for 8 hours. Cell pellets were then collected for analysis. For comparison, we also examined 4T1 cells that were not pre-treated with albumin and were only exposed to BSA@SP1 nanoparticles. The afterglow images revealed that cells not pre-treated with albumin displayed high afterglow luminescence signals, indicating significant SPARC-mediated uptake in 4T1 tumor cells (Fig. S25). In contrast, cells pre-treated with albumin showed saturated SPARC receptors, resulting in lower uptake of BSA@SP1 nanoparticles and consequently lower afterglow luminescence signals.

### 2.5. *Ex Vivo* Porcine Tissue Permeation Profiling using Tumor-Mimicking Phantoms

In evaluating the potential effectiveness of BSA@SP1 in surgical procedures, it is imperative to understand how their afterglow signals vary when tumors are deeply situated within normal tissues. This understanding is crucial for guiding their future clinical applications. To investigate this behavior of BSA@SP1 across various tissue types, we utilized 3D tumor-mimicking phantom models using a methodology previously developed by us ^32,33^ comprising intralipids to simulate fat content (scattering component), hemoglobin to mimic blood vessels (absorption component), and tumor cells labeled with nanoparticles or solely nanoparticles at *in vivo* concentration levels (fluorescence component).^40,41^ It offers distinct advantages over commonly adopted methods, such as utilizing intralipid solutions for simulation^42,43^ because tumor-mimicking phantoms can more accurately replicate the optical properties of real tumors, including their scattering and absorption coefficients. As image-guided surgery relies on detecting NIR optical signals, typically from dye uptake, thus, employing phantoms that closely simulate these optical characteristics can yield more precise and realistic outcomes compared to simulations with intralipid solutions, especially when compared to actual *in vivo* imaging scenarios. Furthermore, these engineered phantoms maintain their stability over time, ensuring consistent experimental outcomes. In contrast, intralipid solutions may exhibit changes in their optical properties due to factors such as temperature fluctuations or particle aggregation, leading to variability in imaging results. Moreover, tumor phantoms can accurately replicate the structural characteristics of tumors, including their dimensions, and morphology. By regulating the concentration of fluorophores within these phantoms or by imparting tumor cells labeled with fluorophores, we can achieve concentrations closer to those observed *in vivo* mouse models.

Here, we aimed to conduct tissue permeation profiling using BSA@SP1 tumor-mimicking phantoms. We evaluated how their afterglow luminescence and fluorescence signals varied when covered with layers of porcine fat, muscle, and skin of varying thicknesses. The BSA@SP1 tumor-mimicking phantoms were prepared (Figure 4a) as described in the methods section. After covering them fat, muscle, and skin tissues of different depths, we imaged them using an IVIS imager under both fluorescence and bioluminescence imaging modes. Our observations revealed reductions in the fluorescence and afterglow tumor phantom-to-normal tissue (T/NT) ratios when various thicknesses of fat, muscle, and skin tissues were placed on top of the BSA@SP1 phantoms. Specifically, the fluorescence T/NT ratios of BSA@SP1 phantoms experienced a significant drop from ∼22 to ∼6 when covered with a 2mm layer of fat tissue (Figure 4c). This reduction further intensified, reaching ∼2 when a 4mm fat tissue, ∼1.5 with 6mm, and eventually dropping to ∼1 with the addition of a 10mm fat tissue addition (Figure 4c). As anticipated, the afterglow T/NT ratios of BSA@SP1 phantoms were higher than fluorescence T/NT ratios due to the former’s background signal compared to fluorescence. The afterglow T/NT ratios of BSA@SP1 decreased from ∼45 to ∼16 when covered with a 2mm layer of fat tissue. This reduction increased progressively, reaching ∼9 when a 4mm fat tissue, ∼6.5 with 6mm, and eventually dropping to ∼5.4 for 8mm fat tissue covering, and 4.5 with the addition of a 10mm fat tissue layer. Remarkably, these T/NT ratios remained above the Rose criterion of 5,^44-46^ even up to 8mm depth, underscoring the promising potential of afterglow luminescence in deep tissue imaging. A similar trend was observed for afterglow vs. fluorescence images for BSA@SP1 phantoms when covered with sequential 2mm tissue slices of porcine muscles where the T/NT ratios (>5) for afterglow were significantly higher than fluorescence for tissue slices of till ∼6mm (Figure 4g). Finally, to investigate the signal penetration for BSA@SPNs on the epidermis, we placed porcine skin on top of BSA@SP1 phantoms. A maximum tissue depth of 4mm, as opposed to 10mm for muscle and fat, was selected for skin to replicate the maximum thickness of the human dermis and epidermis layers. We observed again a significant reduction in fluorescent signals, whereas afterglow signals despite dropping had a substantially higher T/NT ratio when compared to fluorescence signals in skin (Figure 4e). The loss of fluorescence signal through the skin or other tissue types is not a significant concern, especially as most image-guided surgeries are open surgeries. However, having the advantage of deeper penetration provided by afterglow luminescence can enable us to locate the precise location for buried tumor lesions surrounding fat and muscle tissues. We would like to highlight a key difference between our study and recent ones on evaluating the depth penetration of afterglow luminescence. While many seminal studies have predominantly used nanoparticle solutions,^12,47^ we opted for 3D tumor-mimicking phantoms. This decision introduced us to certain challenges, particularly intralipid and hemoglobin, which could potentially dampen afterglow luminescence signals. However, these factors were not encountered in the studies using nanoparticle solutions for depth penetration. Additionally, another contributing factor could be the concentration difference between our samples and those in existing literature.^12,47^ We maintained a final concentration of ∼10 μg/mL of BSA@SPs in our phantoms, whereas studies often utilized concentrations approximately ∼ten-fold higher (∼60-100μg/mL) for their depth penetration studies.^12,47^ This disparity in experimental setups might explain why some studies achieved depth penetrations of approximately 10-30mm for afterglow luminescence, whereas our results ranged from 8-10mm. Overall, our findings demonstrate that by conducting depth penetration studies under more realistic *in vivo* conditions, we consistently observed stronger afterglow luminescence signals at deeper tissue depths in comparison to fluorescence signals.

**Figure 4.**
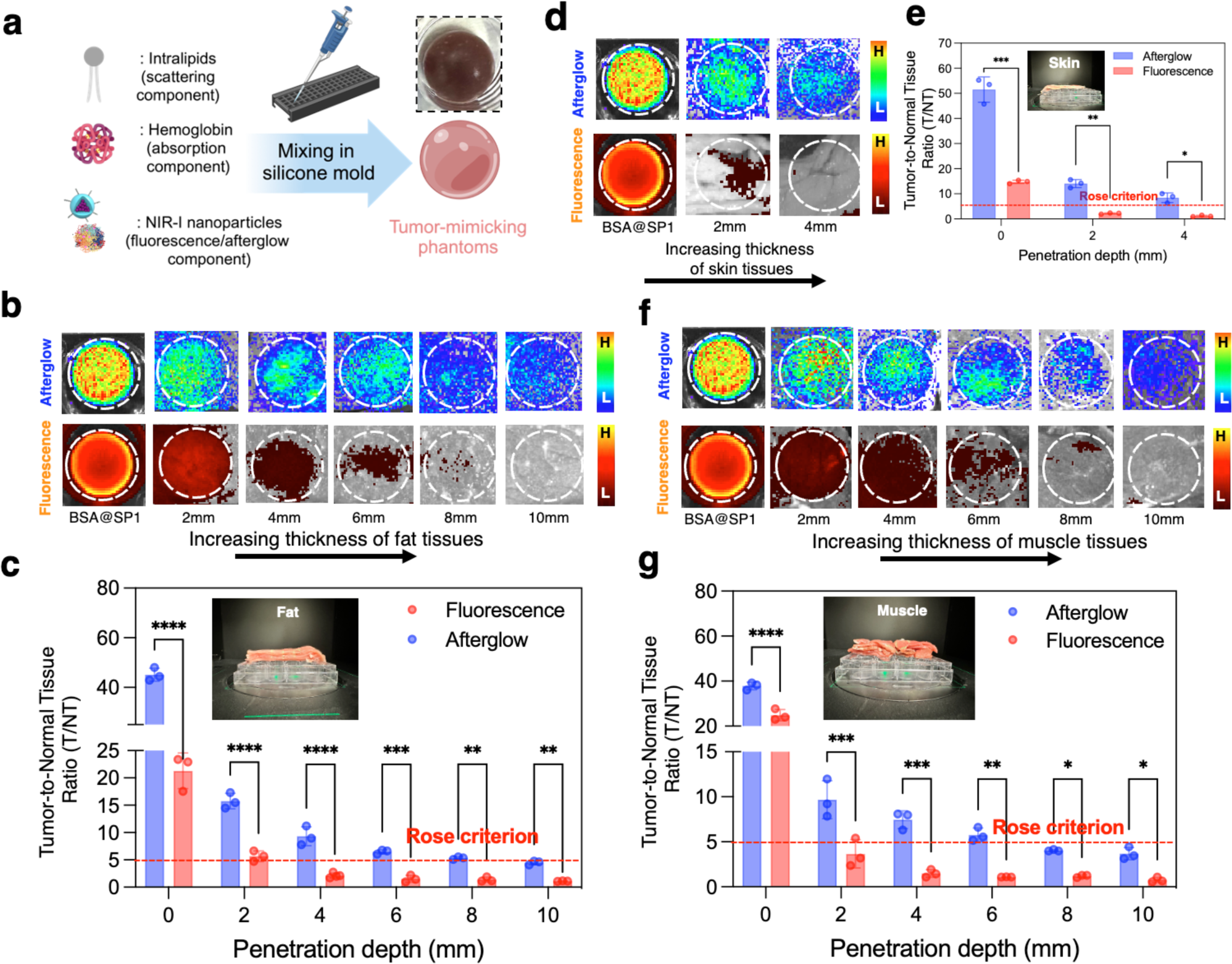
*Ex vivo* imaging of tissue penetration and signal-to-background ratios of BSA@SP1 afterglow luminescence and fluorescence. (a) Schematic process showing the preparation process of tumor-mimicking phantoms and inset showing the picture of a representative as-prepared phantom. Afterglow and fluorescence images of tumor-mimicking phantoms covered with increasing width of tissue slices of (b) fat, (d) skin, and (f) muscle. The corresponding tumor phantom-to-normal tissue (T/NT) ratios for both afterglow and fluorescence were plotted and compared for (c) fat, (e) skin, and (g) muscle.

### 2.6. Afterglow-guided Thoracic Surgery Simulations on *Ex Vivo* Porcine Lungs

Inspired by the superior performance of BSA@SP1 tumor-mimicking phantoms, which exhibit robust T/NT signal ratios in afterglow imaging, we explored their potential applications in afterglow-guided surgery. Our investigation utilized *ex vivo* porcine lungs as experimental models, with strategically implanted tumor-mimicking phantoms at various locations to simulate diverse scenarios akin to image-guided thoracic surgery. Unlike the previous case where we used a solution containing gelatin, intralipid, and hemoglobin to make phantoms, we included tumor cells at a fixed density labeled with BSA@SP1 nanoparticles (Figure 5a). We noticed that decreasing the cell density from 100,000 to 1,000 cells leads to a decrease in the uptake of BSA@SP1 nanoparticles at a concentration of 100 µg/mL, as indicated by the trend observed in their afterglow luminescence signals (Figure S26a) following the creation of the phantoms. Similarly, altering the concentration of BSA@SP1 nanoparticles to 50 µg/mL follows a comparable trend, although the nanoparticles at a concentration of 100 µg/mL exhibit stronger afterglow luminescence signals at cell densities of 100,000 and 10,000 cells (Figure S26b). Two selected locations (Figure 5b) were chosen to simulate different thoracic surgery scenarios: on the surface of the right inferior lobe (location #1), and on the posterior side of the left inferior lobe (location #2). Subsequently, these locations were imaged using both fluorescence and afterglow imaging *via* an IVIS imager. At location #1, characterized by a surface-level tumor phantom placement, we observed strong fluorescence and afterglow signals (Figure 5c). Despite higher background signals from surrounding tissues in fluorescence imaging, the fluorescence signal from the tumor phantom could be discerned (Figure 5c, location #1). The tissue background signal in the afterglow imaging was minimal, allowing only afterglow luminescence signals from the tumor phantoms. In contrast, at location #2, where the tumor phantom was buried deeper within the left inferior lobe, the fluorescence signal diminished significantly, while tissue autofluorescence continued to pose interference (Figure 5d). Conversely, afterglow luminescence imaging at this location showed a strong signal with reduced interferences from surrounding tissues, as anticipated (Figure 5d, location #2). At these two locations, we calculated the T/NT ratios, which were superior in afterglow imaging compared to fluorescence imaging (Figure 5e, f). Moreover, these ratios were significantly higher than the Rose criterion. These findings underscore the potential of afterglow imaging, particularly in scenarios where fluorescence imaging encounters limitations due to tissue autofluorescence and depth of tumor localization. This preliminary *ex vivo* studies further highlights the promising role of afterglow-guided surgery in enhancing surgical precision and outcomes in thoracic procedures. To simulate this process *in vivo*, our BSA@SP1 nanoparticles can be delivered directly into solid tumors via intratumoral injections. Subsequently, light irradiation triggers the rapid generation of afterglow luminescence signals, which can then guide subsequent surgical interventions. This approach holds promise for enhancing the efficiency of the surgical procedure. We wanted to emphasize that employing a specific wavelength laser for afterglow imaging offers notable advantages. Laser light emits precise, narrowband wavelengths, enhancing imaging sensitivity and specificity over white light, thereby improving resolution and accuracy. However, we recognize the challenges posed by the excitation of our BSA@SP1 afterglow probe, which occurs at 605nm, especially in deep tissue scenarios. However, recent advancements have tackled this limitation through innovative strategies involving chemical modulation of these probes. Researchers have been exploring methods to chemically modify these probes specifically for NIR-II imaging,^48^ taking advantage of reduced light scattering, absorption, and autofluorescence. Furthermore, pioneering research in afterglow imaging has highlighted the potential of recharging or cyclic modulation of afterglow probes using ultrasound-induced afterglow (reaching approximately 4cm)^49^ and X-ray-mediated afterglow.^50^ These methods offer deeper penetration depths into tumors, enabling effective excitation of the afterglow probes.

**Figure 5.**
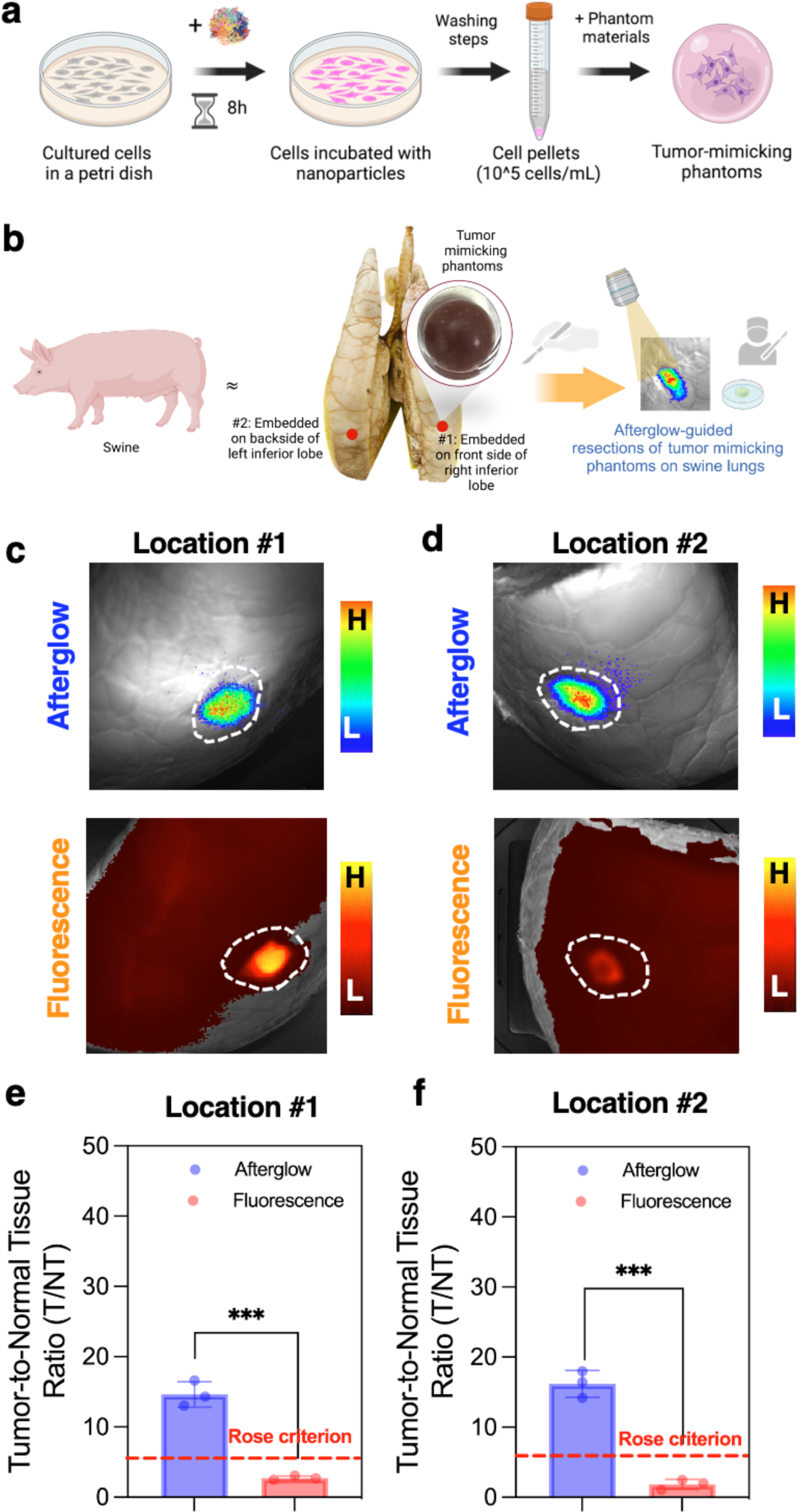
Afterglow luminescence-guided surgery using BSA@SP1 on swine lungs embedded with tumor cell-mimicking phantoms. (a) Schematic representation of preparing tumor-cell mimicking phantoms composed of BSA@SP nanoparticles internalized in 4T1 tumor cells (cell density of ∼100k cells) mixed with phantom solutions. (b) Schematic process showing embedding of tumor cell-mimicking phantoms on *ex vivo* porcine lungs at different locations (#1, #2). (c, d) Afterglow and fluorescence imaging of these two different locations using an IVIS Imager, and (e, f) corresponding T/NT ratios were measured.

### 2.7. Biosafety Evaluation of BSA@SP1 Nanoparticles

Finally, we conducted an erythrocyte hemolysis assay to assess the interactions between BSA@SP1 and porcine-derived red blood cells. As depicted in Figure S27, BSA@SP1 demonstrated minimal hemolysis across all tested concentrations (10–200 μg/mL). These findings collectively suggest that BSA@SPNs exhibit negligible in vivo toxicity to healthy porcine blood, at least within our tested dosage range, making them suitable for potential in vivo applications. Nevertheless, further investigation is warranted to systematically explore the short- and long-term toxicity responses of these nanoparticles across various doses.

## 3. Conclusion

In this report, we have introduced a novel repurposing technique using a BSA coating that enhances the suboptimal afterglow brightness of existing NIR-I afterglow luminescence materials, using a semiconducting polymer (PFODBT/SP1) as a representative example. Through this approach, BSA@SP1 demonstrated a ∼3-fold amplification of the afterglow luminescence signal compared to polymer-lipid-coated PFODBT. Our comprehensive analysis, including molecule docking, physicochemical characterization, and optical assessments, highlights BSA@SP1’s superior afterglow properties, cyclic afterglow behavior, long-term colloidal stability, and biocompatibility. Furthermore, we validated the efficacy of BSA@SP1 in aiding surgical interventions using *ex vivo* tumor-mimicking phantom models. Our porcine tissue permeation profiling studies revealed that afterglow signals, with their high T/NT ratios, outperformed fluorescence signals. We also simulated thoracic surgeries on *ex vivo* porcine lungs with implanted tumor-mimicking phantoms, showcasing the advantages of afterglow luminescence over fluorescence imaging for surgical resections. Thus, our protein-based repurposing strategy holds promise for enhancing the brightness of existing and upcoming NIR-I afterglow materials, making them valuable tools for image-guided surgical interventions.

## 4. Experimental Section

### Materials

All materials were purchased from Sigma-Aldrich and used without any modifications unless stated separately. Pork skin, fat, and muscle tissues were acquired from local grocery stores. Dry-preserved porcine lung was purchased from Nasco Education. The SOSG test kit was received as a complementary gift from Lumiprobe USA.

### Preparation of DSPE-PEG@SP1

For preparing DSPE-PEG@SP1, 0.5 mg of SP1/PFODBT were co-dissolved in 2 mL of THF with 10 mg of surfactant, DSPE-PEG2K. This was followed by a dropwise addition of 8mL PBS1x solution in a 20mL vial under probe sonication for 4 minutes. THF was subsequently evaporated from the solution by stirring at 450 rpm at room temperature overnight. The mixture solution was ultrafiltered at 4500 rpm for 10 minutes and then washed three times with PBS1x to remove any excess precursors. The nanoparticle solution was filtered using a 0.22 micron PTFE (Polytetrafluoroethylene) filter and the final product (DSPE-PEG@SP1) was collected and stored at 4 °C.

### Preparation of BSA@SP1

0.5 mg of SP1/PFODBT were dissolved in 2 mL of THF. 66 mg of BSA was dissolved in 10 mL of PBSx1 in a separate vial and heated to approximately 37°C to ensure homogeneity. SP1 solution was then added dropwise to 8mL of BSA-PBS1x solution and sonicated for 4 minutes. THF was subsequently evaporated overnight by stirring at 450 rpm. The mixed solution was ultrafiltered at 4500 rpm for 10 minutes three times and then washed three times with PBS1x to remove any excess precursors or unreacted BSA. The nanoparticle solution was filtered using a 0.22 micron PTFE (Polytetrafluoroethylene) filter and the final product (BSA@SP1) was collected and stored at 4 °C.

### Preparation of DSPE-PEG@SP2

For preparing DSPE-PEG@SP2, 0.5 mg of SP2/PCPDTBT were co-dissolved in 2 mL of THF with 10 mg of surfactant, DSPE-PEG2K. This was followed by a dropwise addition of 8mL PBS1x solution in a 20mL vial under probe sonication for 4 minutes. THF was subsequently evaporated from the solution by stirring at 450 rpm at room temperature overnight. The mixture solution was ultrafiltered at 4500 rpm for 10 minutes and then washed three times with PBS1x to remove any excess precursors. The nanoparticle solution was filtered using a 0.22 micron PTFE (Polytetrafluoroethylene) filter and the final product (DSPE-PEG@SP2) was collected and stored at 4 °C.

### Preparation of BSA@SP2

0.5 mg of SP2/PCPDTBT was dissolved in 2 mL of THF. 66 mg of BSA was dissolved in 10 mL of PBSx1 in a separate vial and heated to approximately 37°C to ensure homogeneity. SP2 solution was then added dropwise to 8mL of BSA-PBS1x solution and sonicated for 4 minutes. THF was subsequently evaporated overnight by stirring at 450 rpm. The mixed solution was ultrafiltered at 4500 rpm for 10 minutes three times and then washed three times with PBS1x to remove any excess precursors or unreacted BSA. The nanoparticle solution was filtered using a 0.22 micron PTFE (Polytetrafluoroethylene) filter and the final product (BSA@SP2) was collected and stored at 4°C.

### Physicochemical and Optical Characterization

Dynamic light scattering (DLS) measurements were conducted to measure the hydrodynamic diameter of nanoparticles using a Litesizer DLS 700 Particle Analyzer instrument (Anton Paar). For each measurement, 50 µL of nanoparticle solution and 950 µL of PBSx1 were used. ζ-Potential measurements were also taken using the Litesizer DLS 700 Particle Analyzer instrument (Anton Paar). Ultraviolet-visible (UV-VIS) absorbance was recorded on a Genesys 30 Visible Spectrophotometer (ThermoFisher Scientific). The absorption spectra were collected at an interval of 1nm scanning from 325nm to 1100nm. Fluorescence spectra were collected on an RF-6000 Spectro Fluorophotometer (Shimadzu). The emission spectra were collected at an interval of 1nm scanning from 690nm to 800nm.

### Transmission Electron Microscopy (TEM) Imaging

A small amount of the nanoparticle solution containing either DSPE-PEG@SP1 or BSA@SP1 was dropped onto a glow-discharged, carbon-coated TEM grid and incubated for 5-10 minutes. Excess liquid was removed, and samples were left to completely dry overnight in a desiccation chamber. All samples were imaged using a JOEL JEM-2100 Cryo transmission electron microscope at 200 kV.

### Colloidal Stability of BSA@SP1 in a Biological Milieu

The hydrodynamic diameter, PDI, and zeta potential values of BSA@SP1 in PBS1x and 10% plasma solution were measured over time (days 1 - 7). All samples were maintained at 37°C, oscillating in a Solaris 2000 orbital shaker (ThermoFisher Scientific), and after each time point, the hydrodynamic diameter, PDI, and zeta potential values were recorded using Litesizer7000 (Anton Parr).

### Fluorescence and Afterglow Imaging using IVIS Imager

Fluorescence and afterglow imaging of nanoparticles (DSPE-PEG@SP1 or BSA@SP1) were conducted using an IVIS imager. ∼200 μL solutions of either DSPE-PEG@SP1 or BSA@SP1 were prepared in centrifuge tubes for both fluorescence imaging and afterglow imaging. For fluorescence imaging, the fluorescence was acquired at an excitation wavelength of 605nm, and emission wavelength of 720nm. For afterglow imaging, the samples were first irradiated for different durations using a 1000-lumen LED white-light source. Subsequently, they were imaged in the IVIS imager under the open filter setting and bioluminescence imaging mode.

### Molecular Docking Calculations

The 3D structure of Bovine Serum Albumin (BSA) (PDB ID: 4F5S) was chosen as the receptor for molecular docking, employing UCF Chimera software. The receptor input file and the ligand structure of PFODBT were prepared separately using Avogadro software, saving them in PDB format for docking. Additionally, specific parameters for the active pocket were defined for the grid box in vina1.2.3 program: x=42.35, y=32.77, z=98.61, with dimensions x=131.07, y=83.35, z=117.02. Default settings were applied for exhaustiveness=8, number of modes=9, and energy range=3 during docking. Post-generation of the requisite files, default parameters were retained for the PFODBT molecule. Subsequently, the best scoring conformation (with an energy of -6.1 kcal/mol), was identified and merged with the protein, saved as a complex, and subjected to analysis using Pymol to elucidate the small molecule-receptor mode of action. In Pymol software, the final pdbqt file of the ligand was combined with the receptor’s final pdb file. Employing the ‘ligand site’ preset mode, all amino acid residues interacting with the ligand were selected and labeled. Furthermore, utilizing the ‘measurement wizard’, the distances between the ligand and receptor amino acids were calculated in Angstrom units.

### Singlet Oxygen Detection

Singlet oxygen sensor green (SOSG) was used to evaluate the singlet oxygen generation by PFODBT/SP1 in DSPE-PEG@SP1 and BSA@SP1 after light irradiation. Both nanoparticles (∼10 μg/mL) were mixed with SOSG (final concentration of 5 μM) and irradiated with white light for a series of different times. Subsequently, the fluorescence intensity of SOSG was measured at 505nm of excitation and the singlet oxygen generation was quantified by comparing the SOSG fluorescence emission at 525nm.^51^

### MTT Assay of DSPE-PEG@SP1 and BSA@SP1

The cytotoxic effect of the nanoparticles in 4T1 cells was investigated using an MTT assay established previously.^52,53^ *In Vitro Cellular Uptake Experiments.* Flow cytometry was conducted to quantify cellular uptake. 4T1 cancer cells were incubated with DSPE-PEG@SP1 and BSA@SP1 at a concentration of 50 μg/mL for 8 hours. Subsequently, the treated cells were washed with PBS 1x, trypsinized, collected, and resuspended in a 1% FBS in PBS solution for cellular uptake analysis using a BD LSRFortessa.

### Tumor Cell-Mimicking Phantom Models Preparation

Tumor cell-mimicking phantoms were prepared using a silicone mold. These phantoms consist of a mixture containing 720 mg type-A gelatin, 150 mg human hemoglobin, 900 µL of intralipid 20%, and 17.1 mL of PBS1x. Firstly, the gelatin is activated in the PBS1x at 60°C with stirring at 950 rpm for 30 minutes. Once the gelatin is activated, the biological components are added to the solution. The vial is inverted, vortexed, and mixed at 60°C with stirring at 950 rpm for an additional 10 minutes. Next, the mold wells are filled with 500 µL of either 50 µg/mL of DSPE-PEG@PFODBT or BSA@PFODBT and 1.5 mL of the phantom stock solution. For the control phantom, 500 µL of PBS1x is used instead of either DSPE-PEG@PFODBT or BSA@PFODBT solution. The mold is placed in the freezer, and the tumor phantoms are allowed to solidify for a minimum of one hour. Each tumor phantom created using the mold has a uniform total volume of 2 mL for every replicate batch.

### Ex Vivo Porcine Tissue Penetration Experiments

To replicate the properties of human skin, muscle, and fat tissues, three distinct porcine tissue samples were utilized. Skin tissue was simulated using a slice of pork belly, chosen for its similarity in texture and composition. Pork tenderloin, known for its low-fat content relative to other swine tissues, served as simulated muscle tissues. Fat (adipose) tissue was simulated using a fatty cut of pork chop. All tissues were procured from local supermarkets and uniformly sliced to a thickness of 2mm using a deli meat slicer (Anescva Premium Slicer Machine) in the laboratory.

To accommodate for the variability in tumor depth, the tissue slices were stacked accordingly. Skin tissues were stacked up to a depth of 4 mm, representing the maximum thickness observed in human skin. Muscle and fat tissues were stacked at various depths at which tumors may occur, ranging from surface-level to buried tumor lesions. Tumor-mimicking phantoms were prepared using a solution of 720 mg type A gelatin, 17.1 mL of PBS1x, 150 mg human hemoglobin, and 900 µL of intralipid 20%. In a silicone mold, a combination of 500 µL of 1X PBS (control) and either 500 µL of 50 µg/mL DSPE-PEG@PFODT or BSA@PFODBT were added, along with 1.5 mL of phantom stock solution, was used to create the tumor-mimicking phantoms for imaging. These phantoms were then placed in a 12-well plate for further analysis. Fluorescence imaging was conducted in the IVIS imager, at an excitation filter of 605 nm and an emission filter of 720 nm for both DSPE-PEG@PFODBT and BSA@PFODBT. For afterglow imaging, before the addition of each layer of tissue, all tumor phantoms were irradiated for 60 seconds with a 1000-lumen LED white-light source.

### Afterglow-Guided Surgery on Ex Vivo Porcine Lung

4T1 cancer cells were grown in T-25 flasks until they reached 80% confluency. Then, they were treated with DMEM containing BSA@SP1 nanoparticles at concentrations of either 100 μg/mL or 50 μg/mL for 24 hours. Afterward, following a series of washing steps, the cells labeled with nanoparticles were gathered, counted, and adjusted to the desired cell density through dilution. They were then mixed with the phantom solutions to create phantoms mimicking tumor cells. The phantoms were then embedded in the two selected locations of the lung: (i) embedded 0.5 mm into the ventral surface of the right inferior lobe, (ii) embedded 5mm below the ventral surface of the right inferior lobe. Before each round of imaging using the IVIS in the bioluminescence setting (for afterglow), the tumor phantoms underwent 60 seconds of irradiation with a 1000-lumen LED white-light source held 30 cm from the samples for optimal lumen intensity, utilizing a COAST® Products light source (Model = PX15R), powered by a 3.7V battery rated for 2600 mAh. For fluorescence imaging, the excitation wavelength of BSA@PFODBT was selected as 605 nm and the emission wavelength was selected as 720 nm.

### Hemolysis Assay

We prepared samples of BSA@SP1 at various concentrations ranging from 10 to 200 μg/mL, which were then mixed with an equal volume of a 2% (v/v) solution of red blood cells (RBCs). Deionized water served as the positive control, while normal saline served as the negative control. The mixing solutions were incubated at 37°C for 4 hours. Subsequently, the solutions were centrifuged, and photographs were taken. The supernatants were analyzed using UV–Vis spectroscopy at a wavelength of 542 nm. The percentage of hemolysis (%) was calculated as follows:

% Hemolysis = (A_sample_ – A_negative_) / (A_positive_ - A_negative_) x 100

where A_s_, A_negative_ and A_positive_ is the absorption value of test, negative and positive group, respectively

### Statistical Analysis

GraphPad Prism was used for statistical analysis throughout this work. Statistical comparisons were made by unpaired Student’s *t*-test (between two groups) and one-way ANOVA (for multiple comparisons) with Tukey’s post-hoc analysis. *P* value < 0.05 was considered statistically significant. Quantitative data were shown as mean ± standard deviation (SD).

## Supporting Information

Supporting Information is available from the Wiley Online Library or from the author.

## Author Contributions

I.S. conceived the idea and designed all the experiments. N.B., K.N., I.V., and I.S. performed all the experiments. A.H. performed molecular docking studies. I.S. and M.C. performed *in vitro* experiments. I.S. wrote the first draft of the manuscript with edits from all the authors. All authors have approved the final version of the manuscript. ‡ N.B. and K.N. contributed equally.

## Acknowledgements

I.S. acknowledges the Edward E. Whitacre Jr. College of Engineering and Texas Tech University for research support. All graphics in this paper were made in Biorender.

Received: ((will be filled in by the editorial staff))

Revised: ((will be filled in by the editorial staff))

Published online: ((will be filled in by the editorial staff))

---

A new approach has been developed to enhance the afterglow luminescence of existing substrates by using a functional protein (bovine serum albumin) coating. Its application in surgical navigation and image-guided cancer surgery has been demonstrated with tumor-mimicking phantoms and ex vivo porcine models.

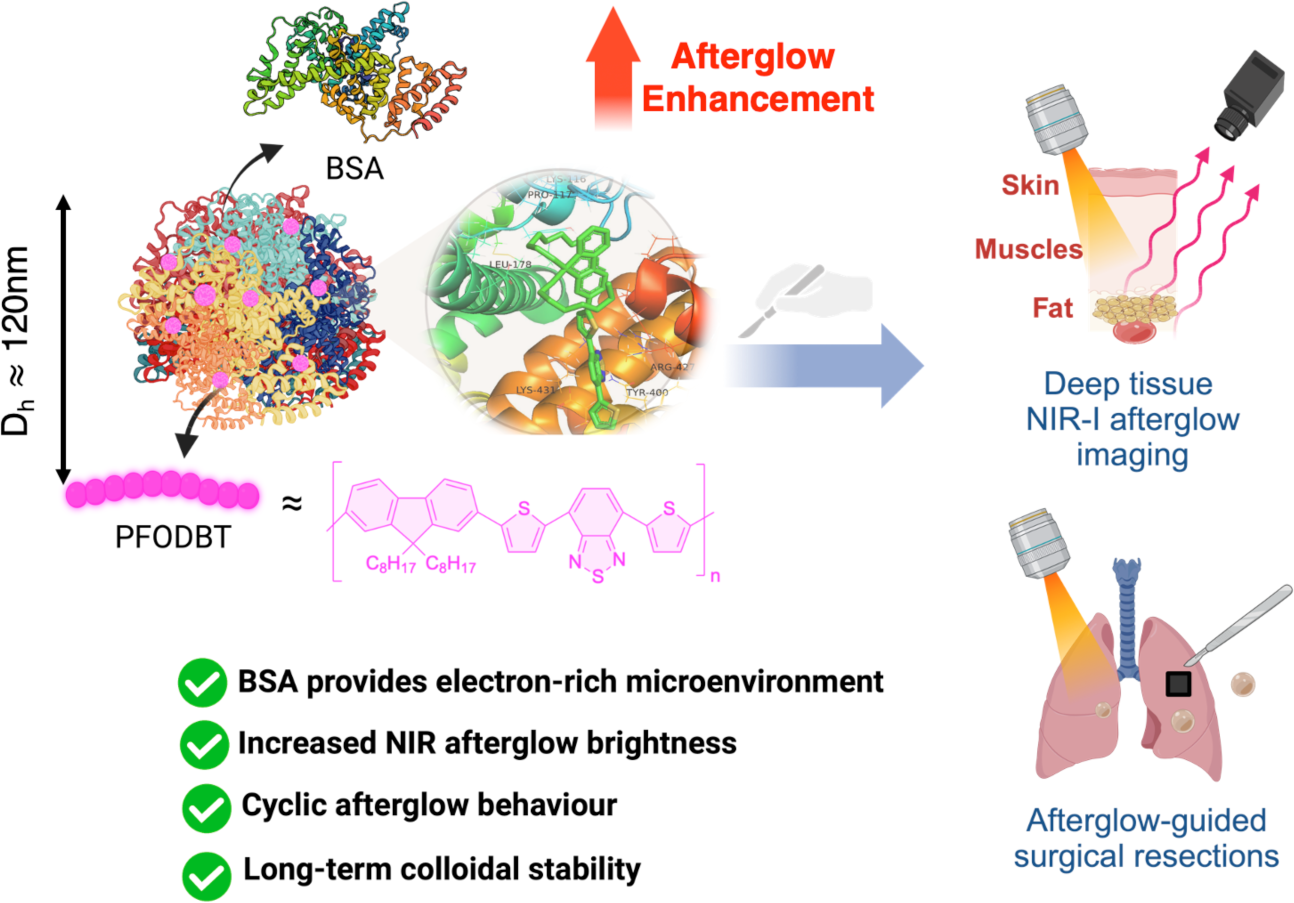

## Supporting Information

**Figure S1.**
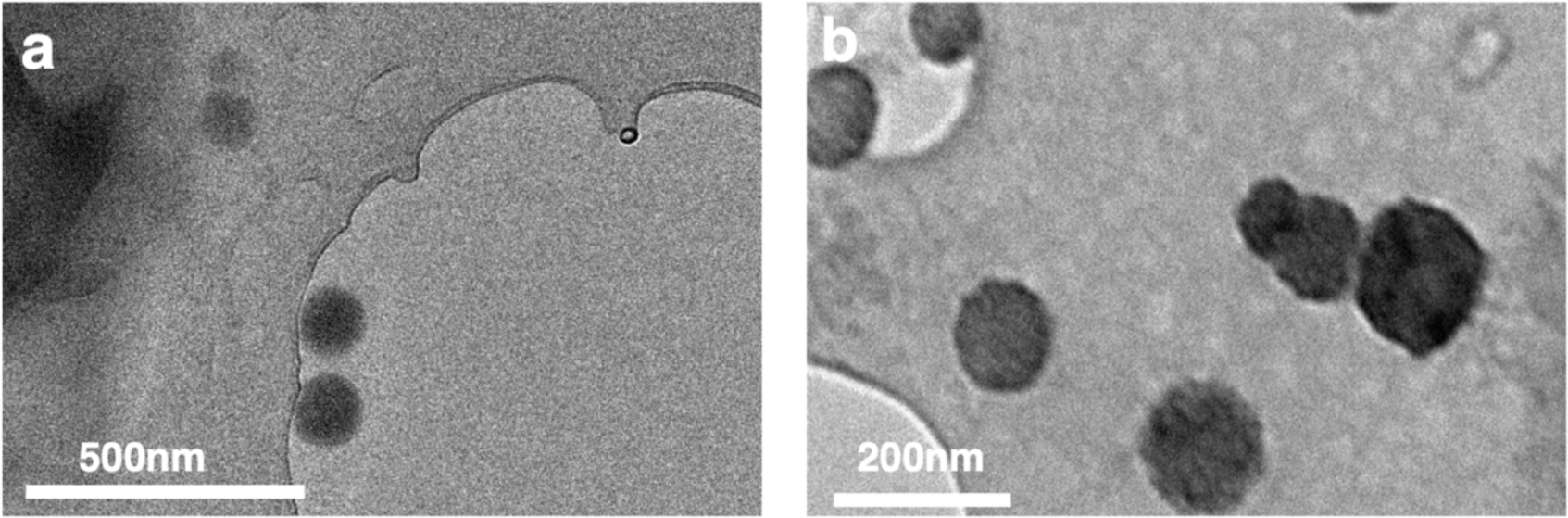
Representative transmission electron microscopy (TEM) images of (a) DSPE-PEG@SP1, and (b) BSA@SP1.

**Figure S2.**
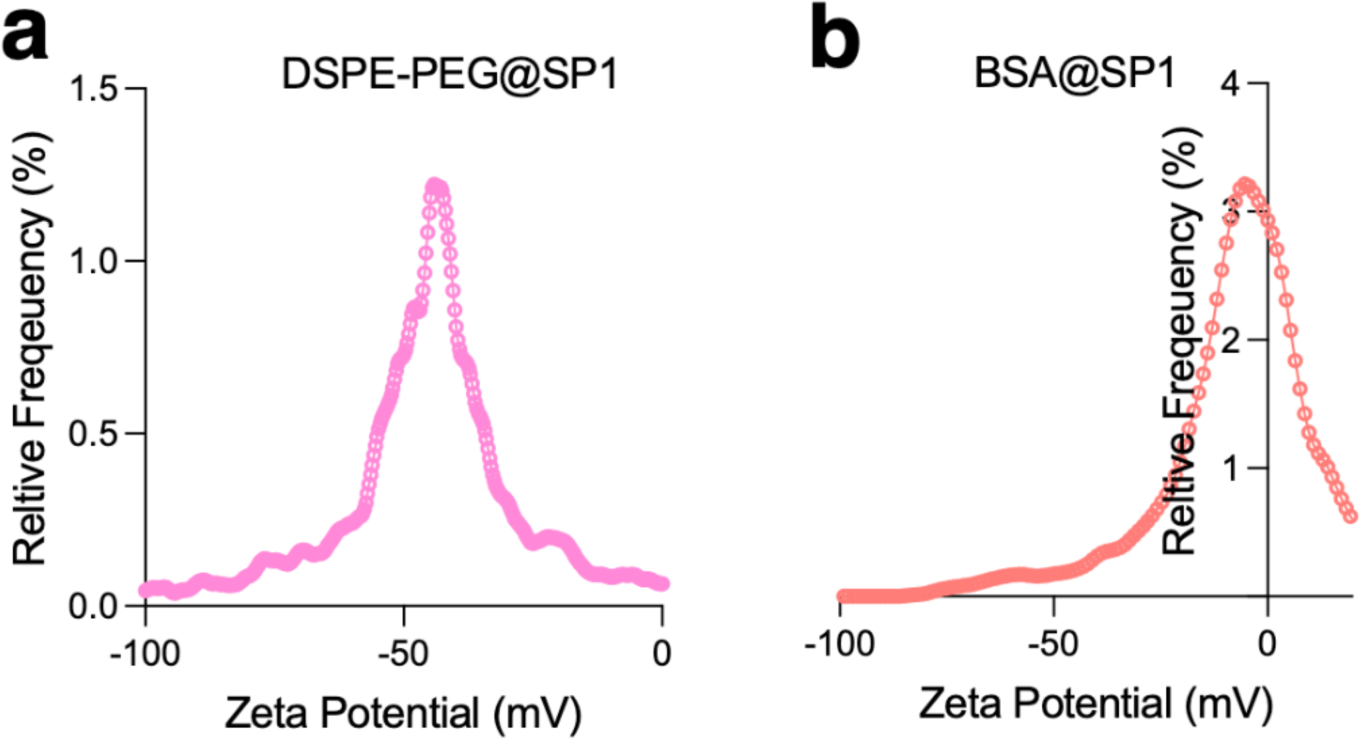
Zeta potential distributions for (a) DSPE-PEG@SP1 and (b) BSA@SP1.

**Figure S3.**
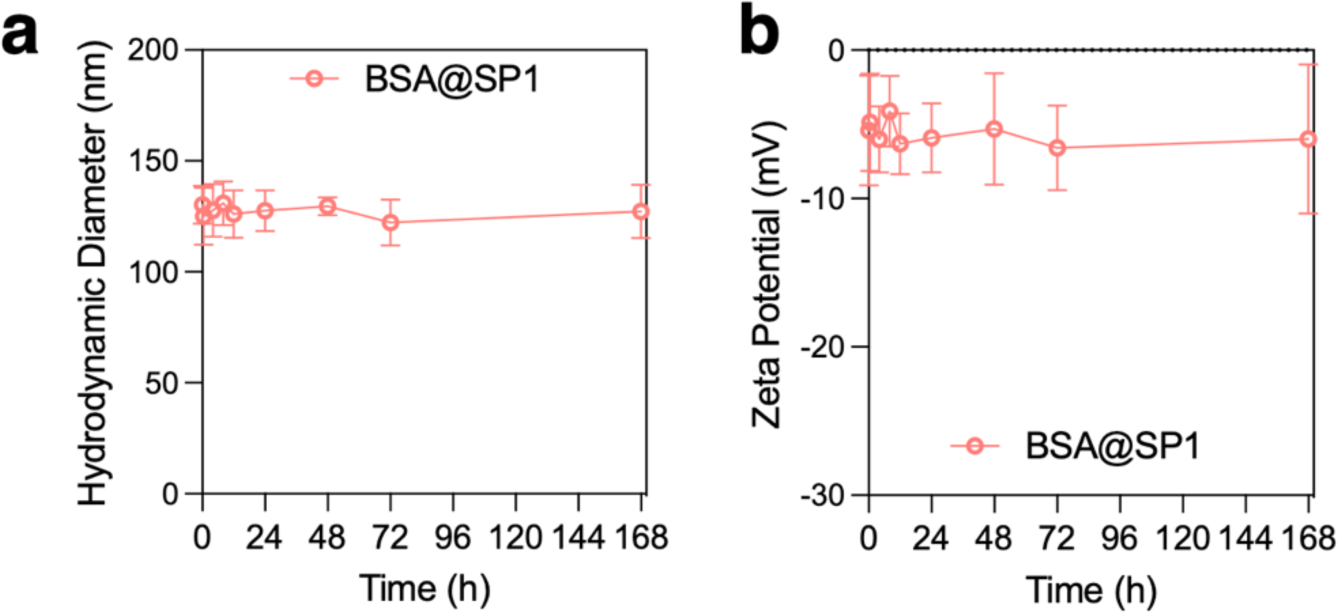
Long-term colloidal stability measurements (∼1 week) for BSA@SP1 in terms of their hydrodynamic diameter and zeta potential measurements.

**Figure S4.**
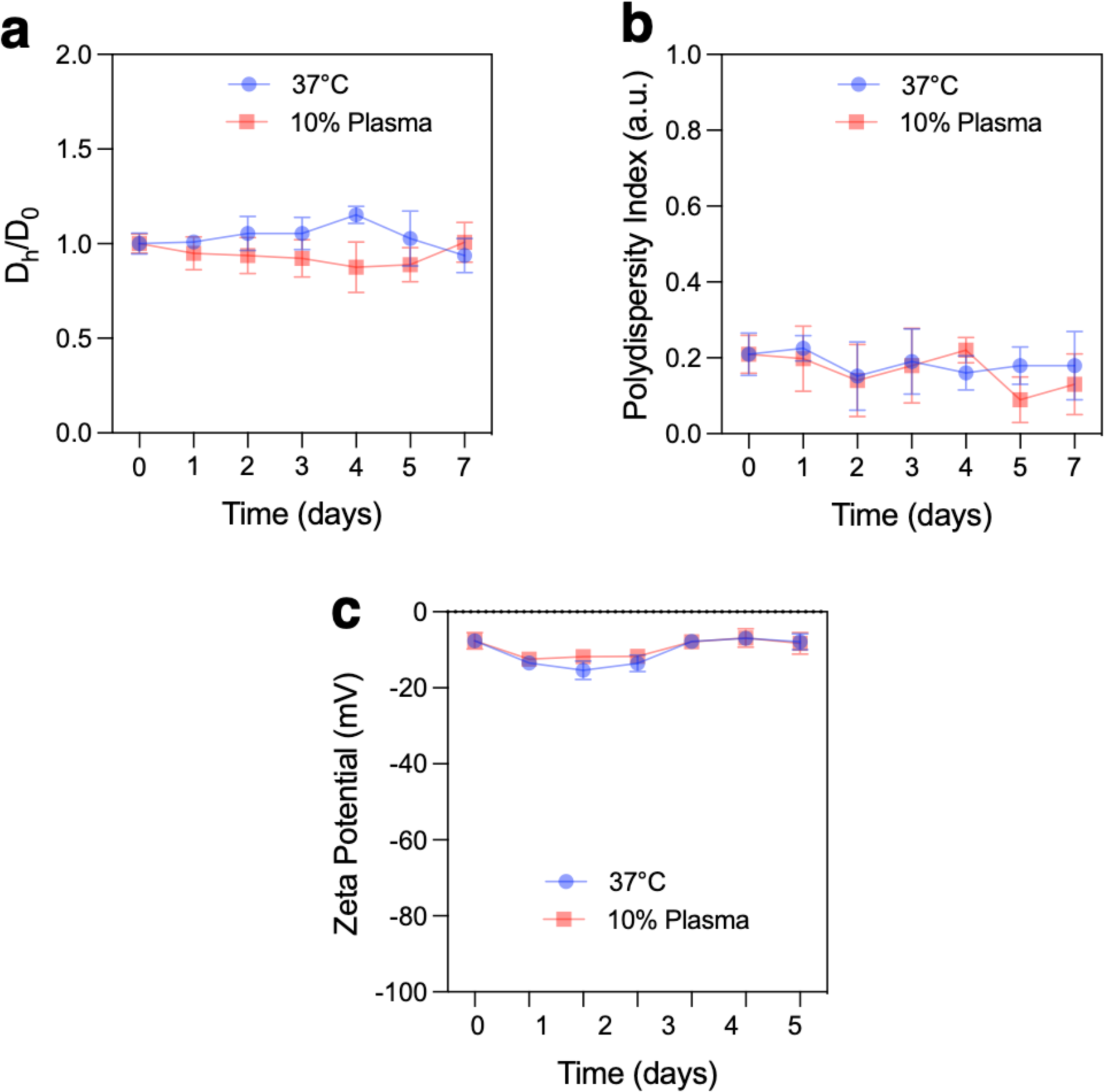
Long-term colloidal stability measurements (∼1 week) for BSA@SP1 in physiological temperature (37°C) and 10% plasma regarding their hydrodynamic diameter and zeta potential measurements.

**Figure S5.**
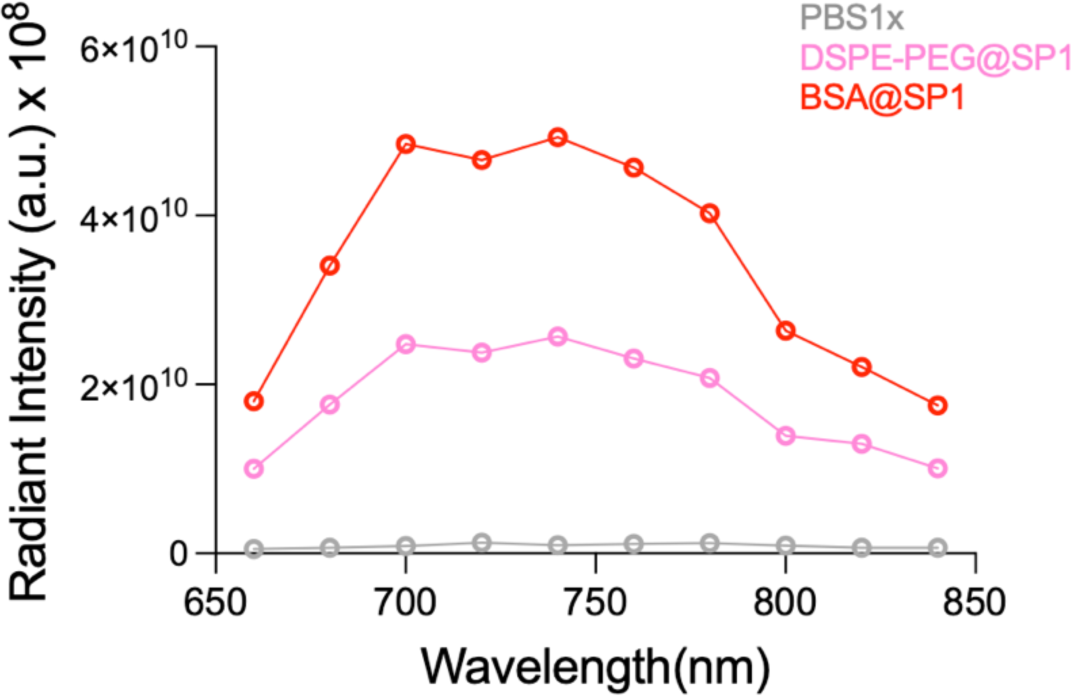
Fluorescent emission spectra obtained from IVIS imager at 605nm excitation for PBS1x, DSPE-PEG@SP1, and BSA@SP1.

**Figure S6.**
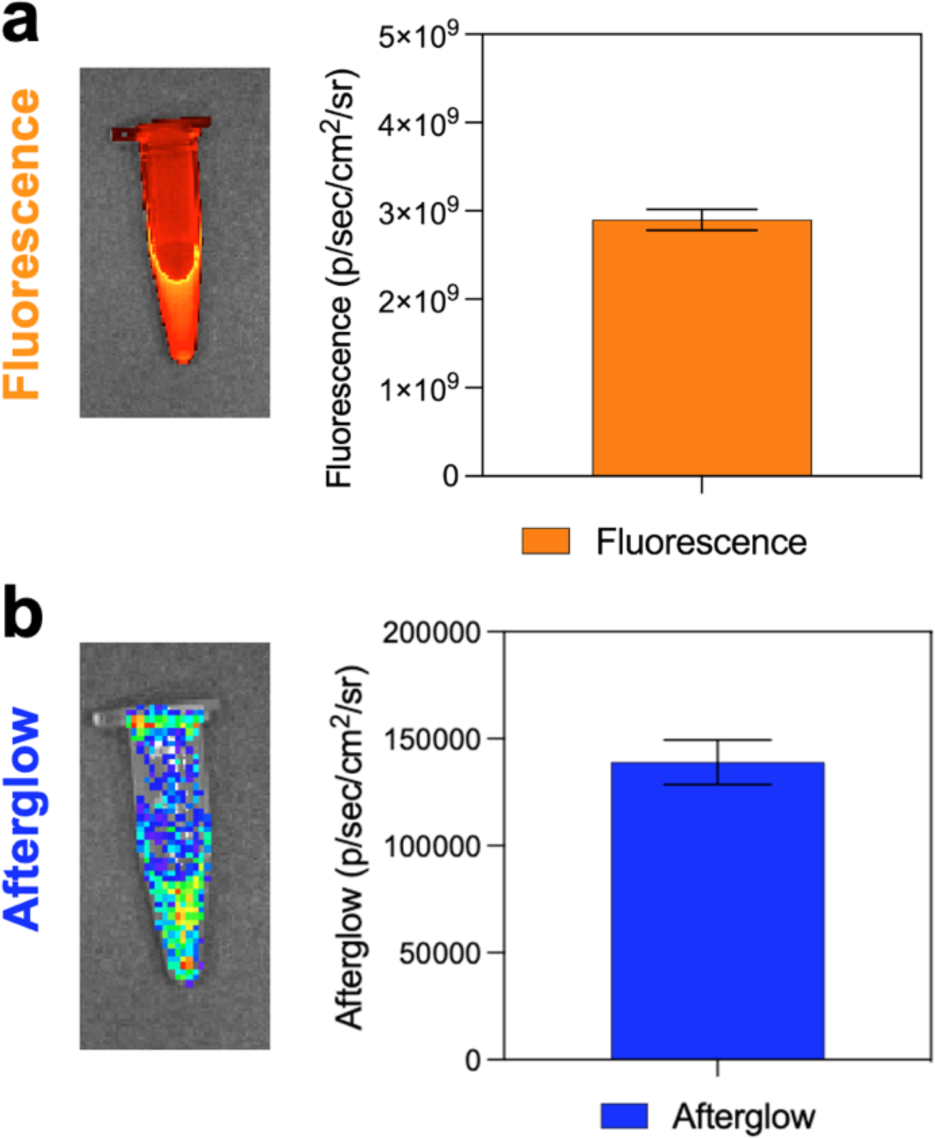
(a) Fluorescence images of SP1/PFODBT taken with an IVIS Imager at an excitation wavelength of 605 nm and an emission wavelength of 720 nm. (b) Afterglow images of SP1/PFODBT captured using the bioluminescence mode with an open filter.

**Figure S7.**
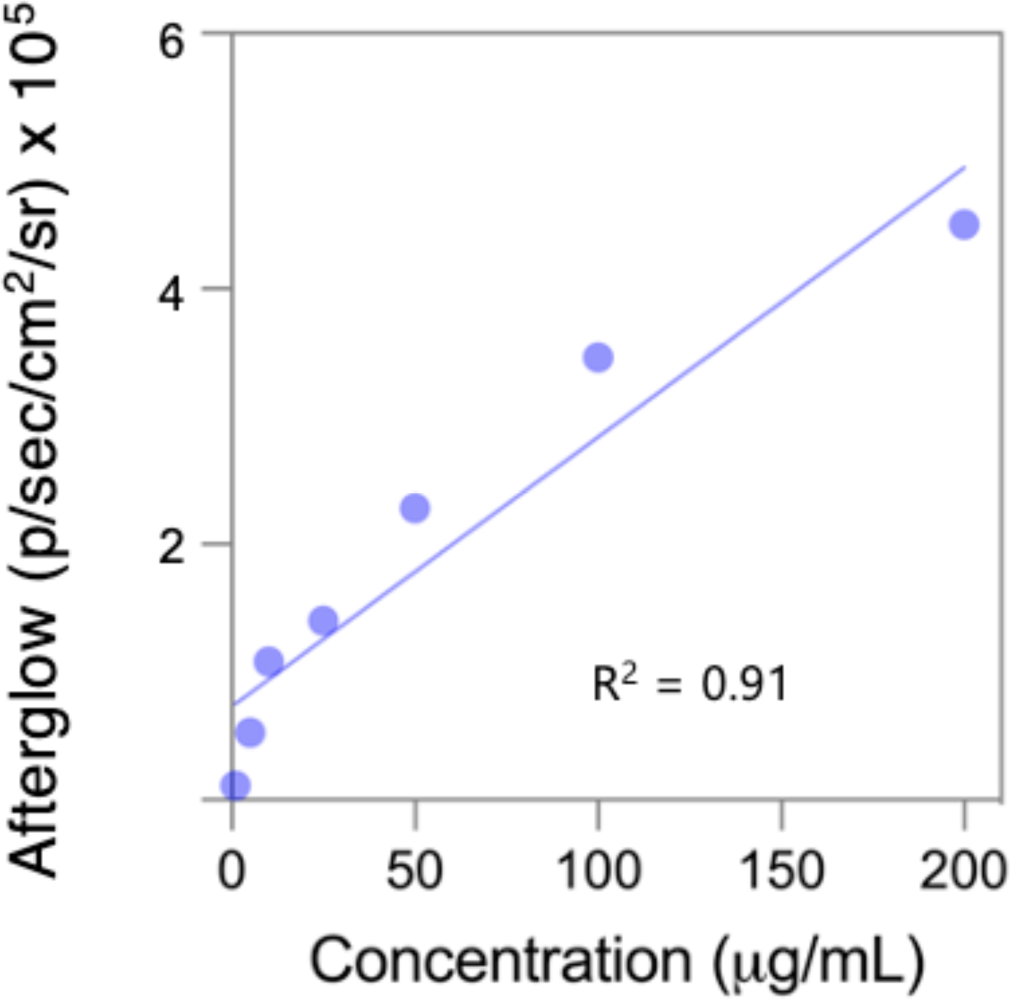
The concentration of BSA@SP1 nanoparticles was varied, and the corresponding afterglow luminescence intensities were measured using an IVIS imager in bioluminescence mode with an open filter. A linear relationship was observed through fitting analysis, with an R² value of 0.91.

**Figure S8.**
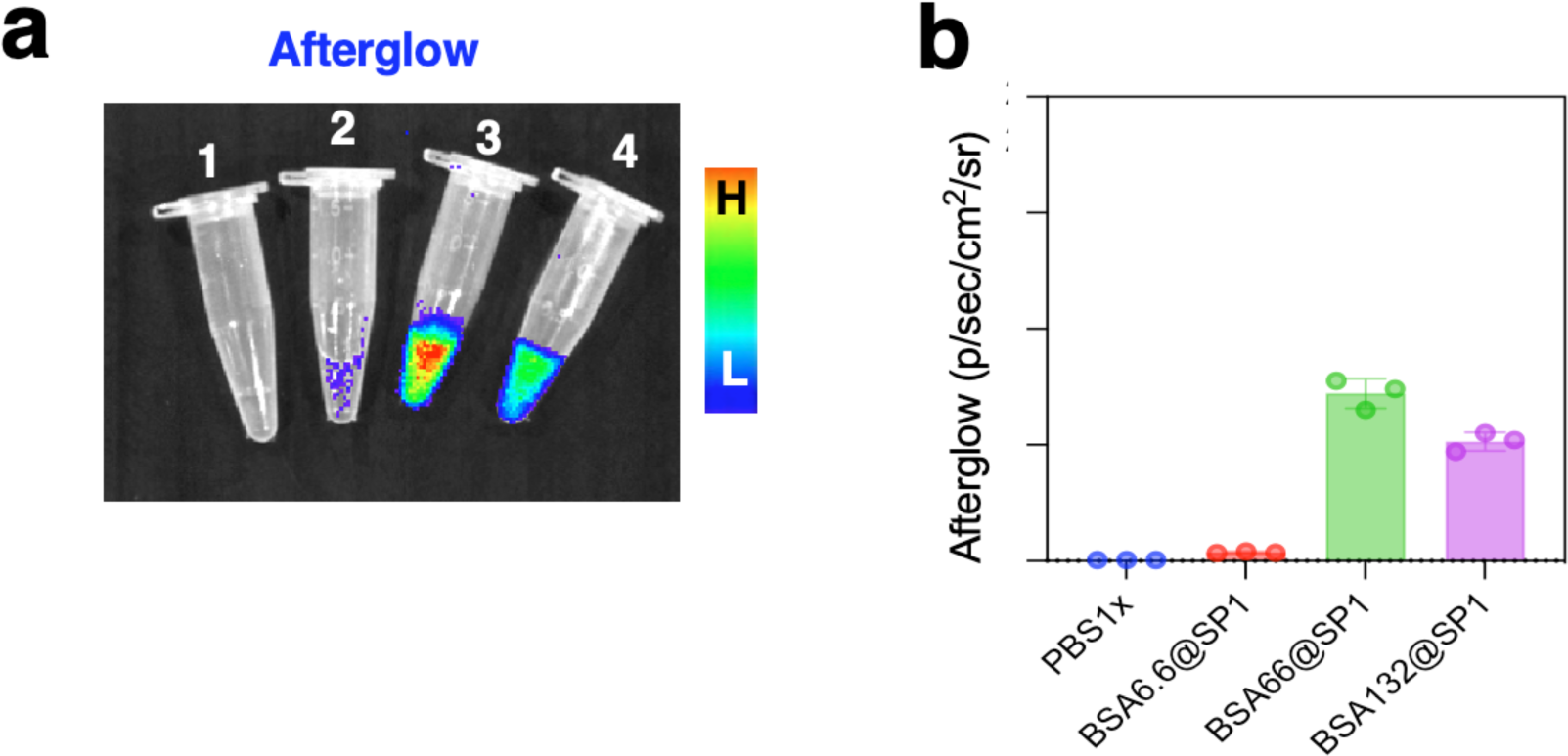
Various concentrations of BSA (6.6 mg, 66 mg, 132 mg each in 10mL PBS1x) were utilized in the nanoprecipitation process to produce BSA@SP1, denoted as BSA6.6@SP1, BSA66@SP1, and BSA132@SP1, respectively. (a) Afterglow images of samples (1) PBS 1x, (2) BSA6.6@SP1, (3) BSA66@SP1, and (4) BSA132@SP1 were captured under open excitation, and their corresponding intensities were measured.

**Figure S9.**
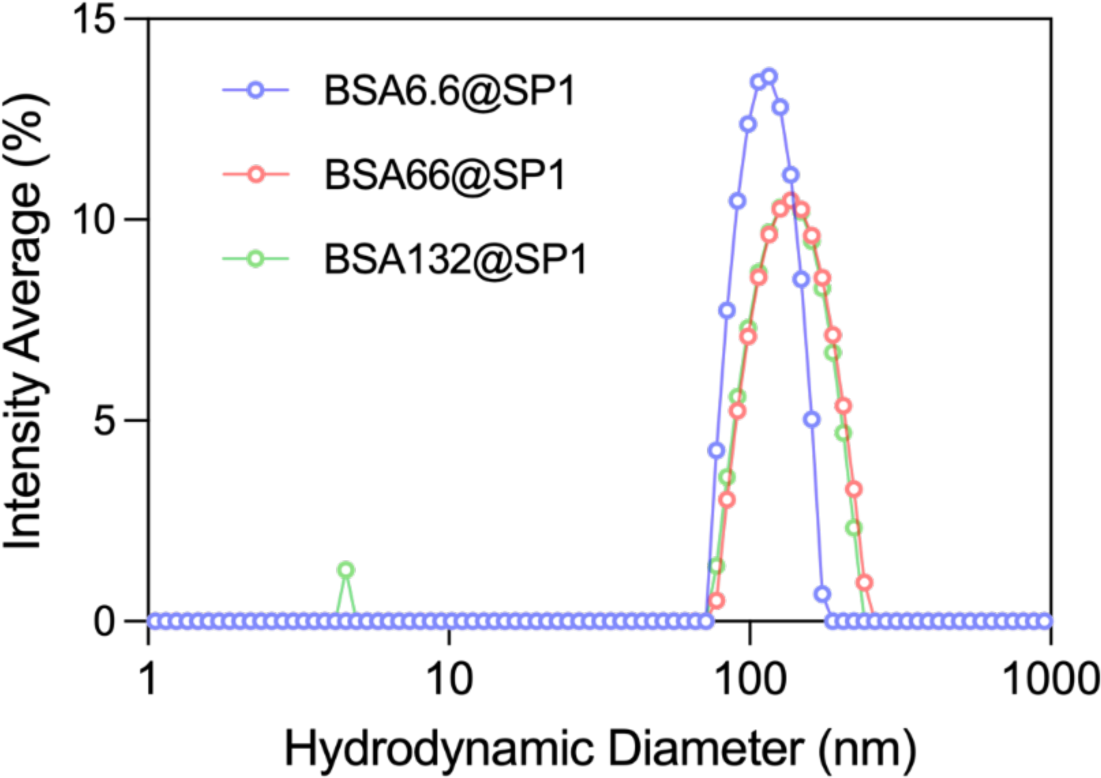
Various concentrations of BSA (6.6 mg, 66 mg, 132 mg) were utilized in the nanoprecipitation process to produce BSA@SP1, denoted as BSA6.6@SP1, BSA66@SP1, and BSA132@SP1, respectively. Hydrodynamic diameter measurements of samples were recorded.

**Figure S10.**
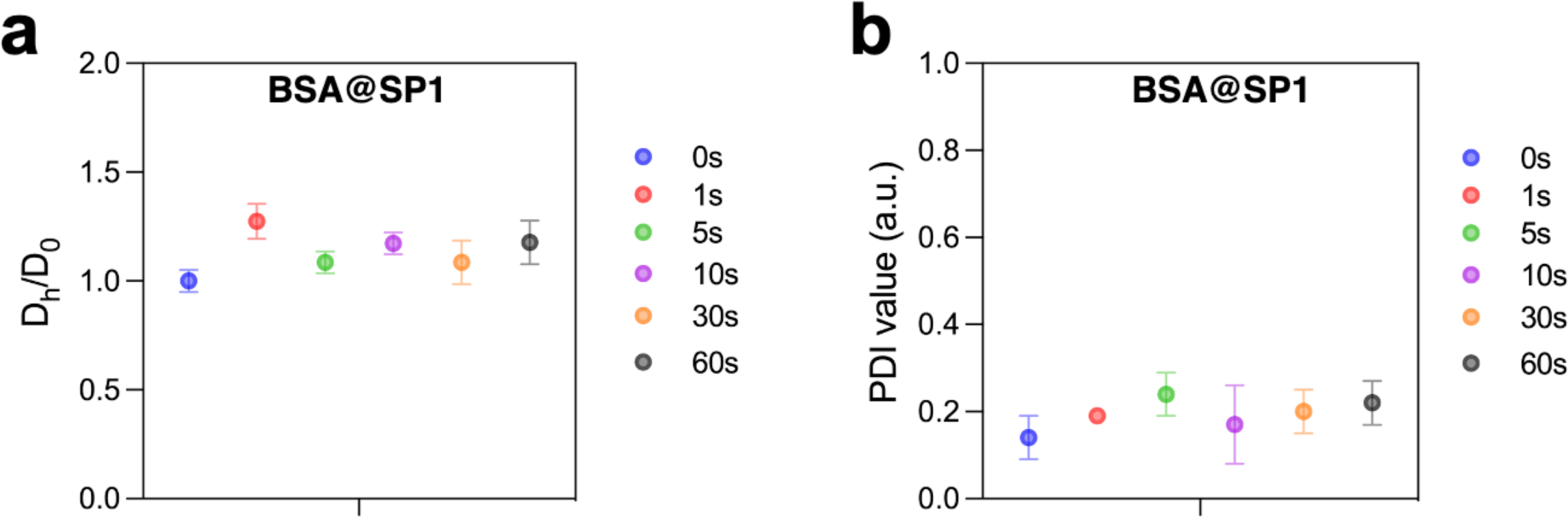
(a) Ratios of hydrodynamic diameters before (D_o_) and after (D_h_) white light irradiation treatment, and (b) corresponding polydispersity values as a function of white light irradiation time.

**Figure S11.**
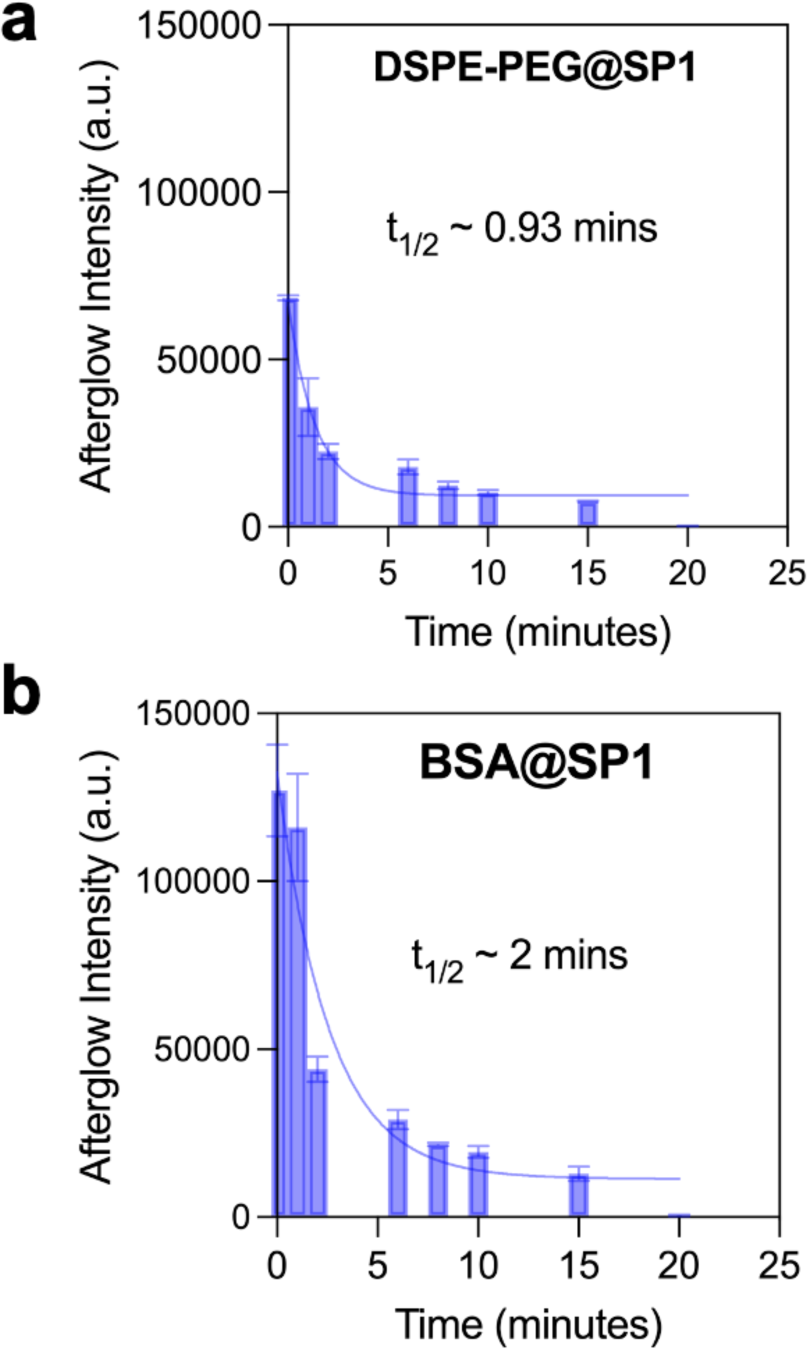
Afterglow lifetime measurements were conducted for (a) DSPE-PEG@SP1 and BSA@SP1 nanoparticles and modeled using a one-phase decay.

**Figure S12.**
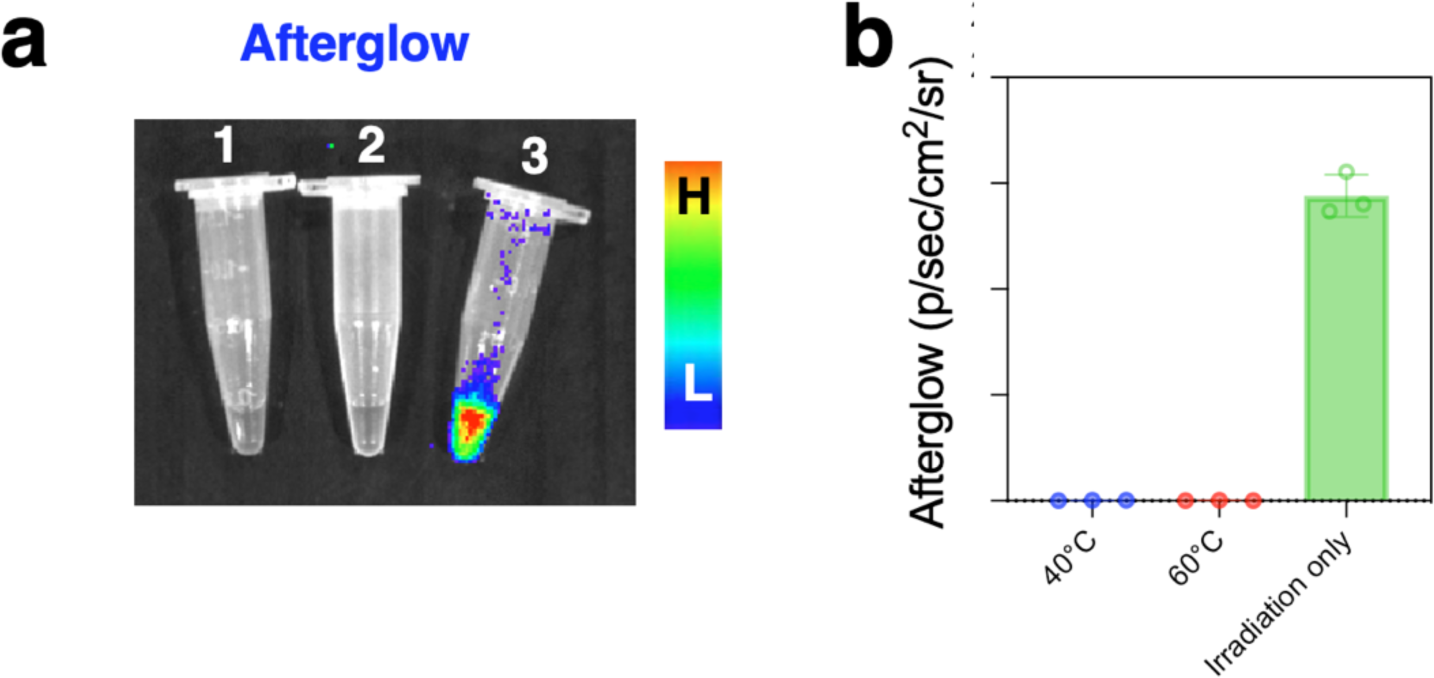
(a) Afterglow images of BSA@SP1 excited by heating at 40°C (1) and 60°C (2), or (3) white light irradiation for 60 seconds. (b) Corresponding quantification of afterglow intensities of the samples.

**Figure S13.**
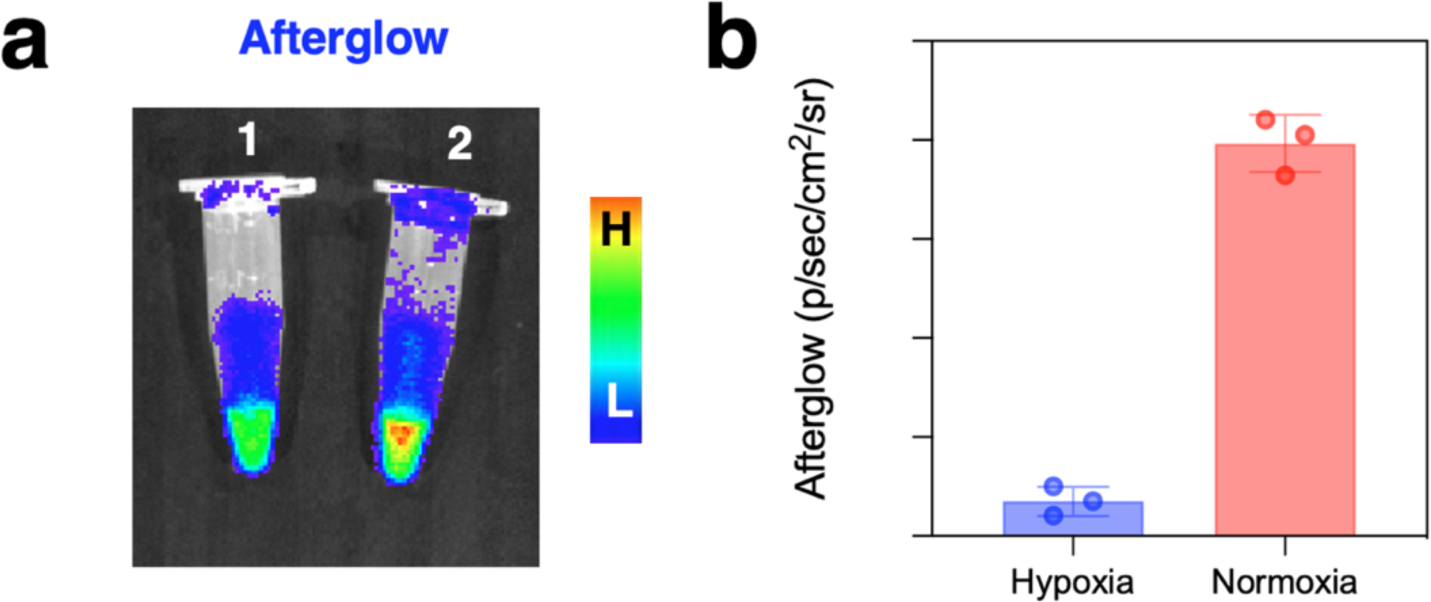
(a) Afterglow emission images of BSA@SP1 under (1) hypoxic, and (2) normoxic conditions after 1min irradiation. (b) Corresponding afterglow intensities were quantified.

**Figure S14.**
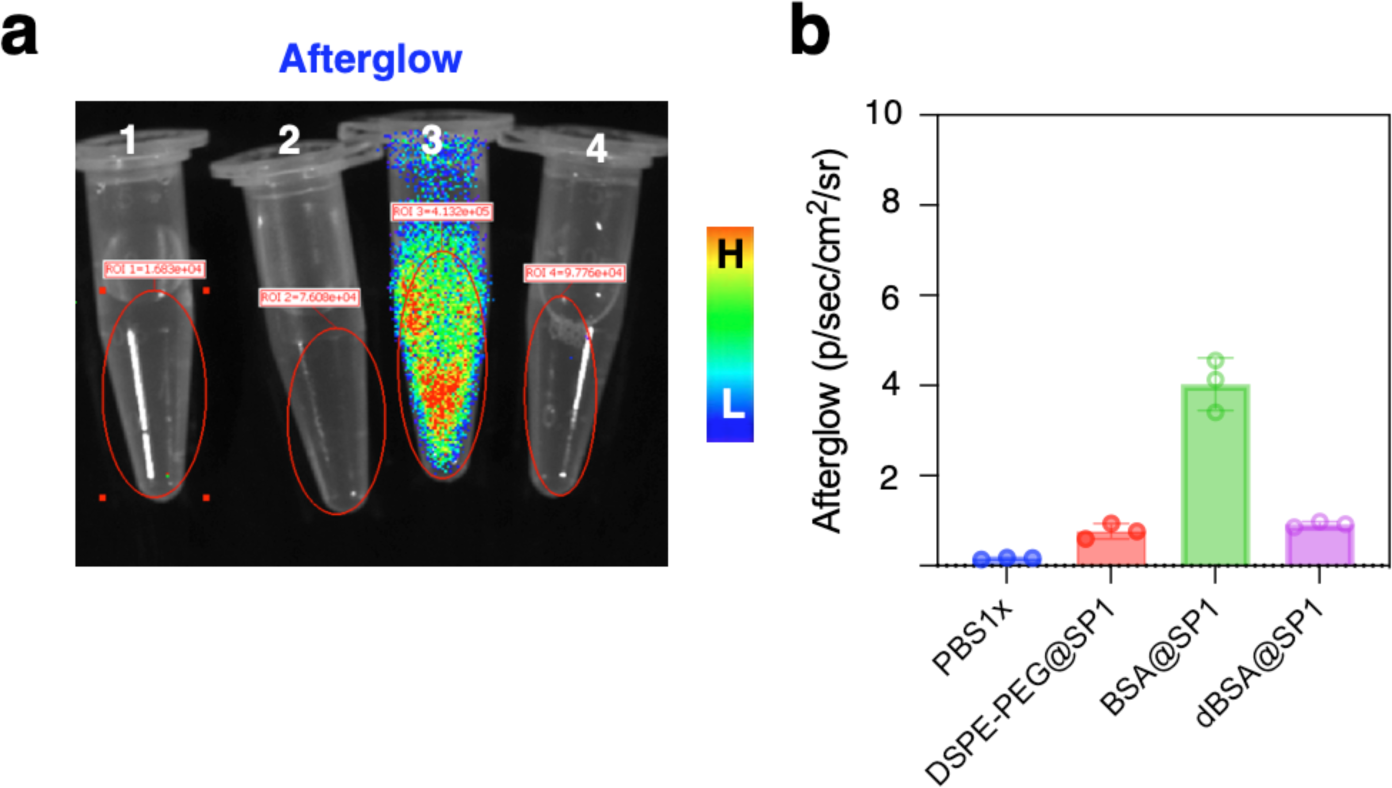
(a) Afterglow images of (1) PBS1x, (2) DSPE-PEG@SP1, (3) BSA@SP1, and (4) denatured BSA@SP1, and (b) their corresponding afterglow luminescence intensities were collected.

**Figure S15.**
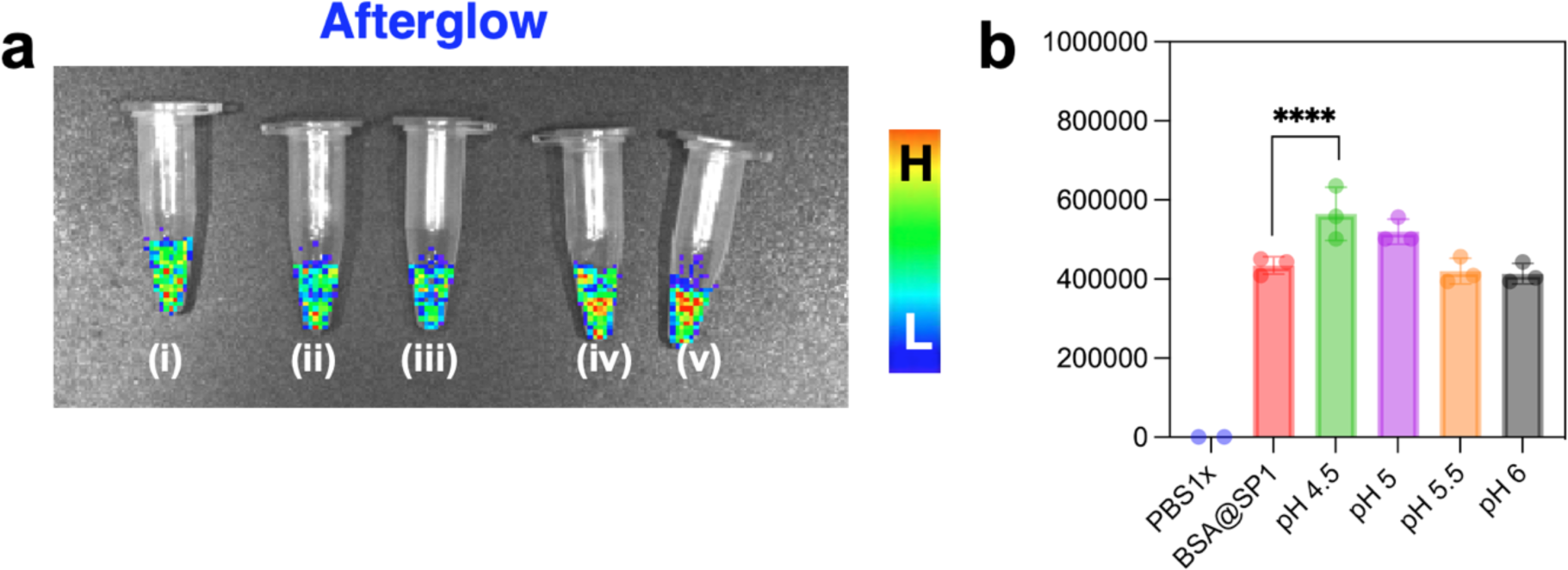
(a) Afterglow images of BSA@SP1 in (i) PBS1x, (ii) pH 4.5, (iii) pH 5, (iv) pH 5.5, and (v) pH 6, and (b) their corresponding afterglow luminescence intensities were collected.

**Figure S16.**
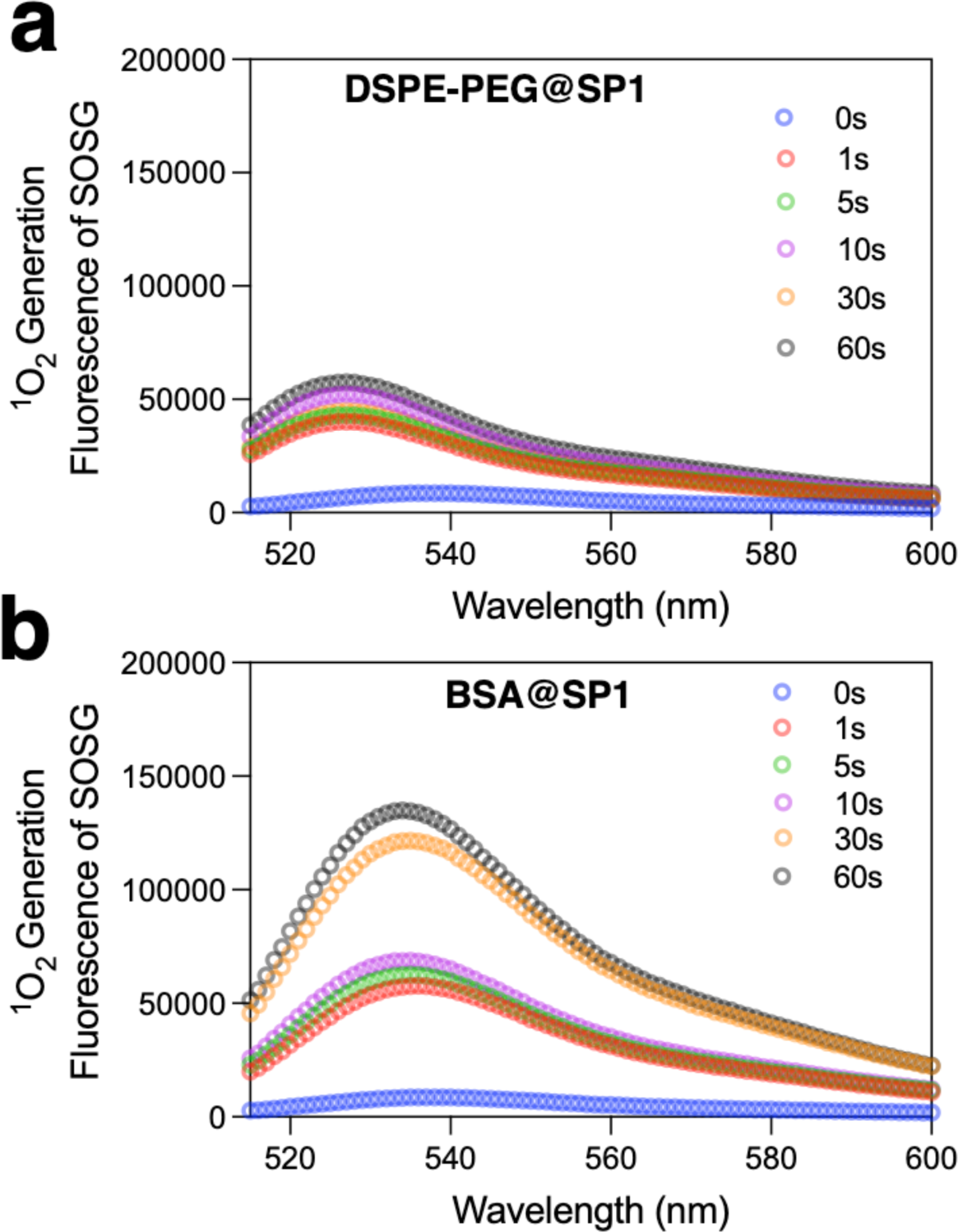
SOSG fluorescent spectra for (a) DSPE-PEG@SP1 and (b) BSA@SP1 correlated with their ^1^O_2_ generation treated with irradiations for different time durations.

**Figure S17.**
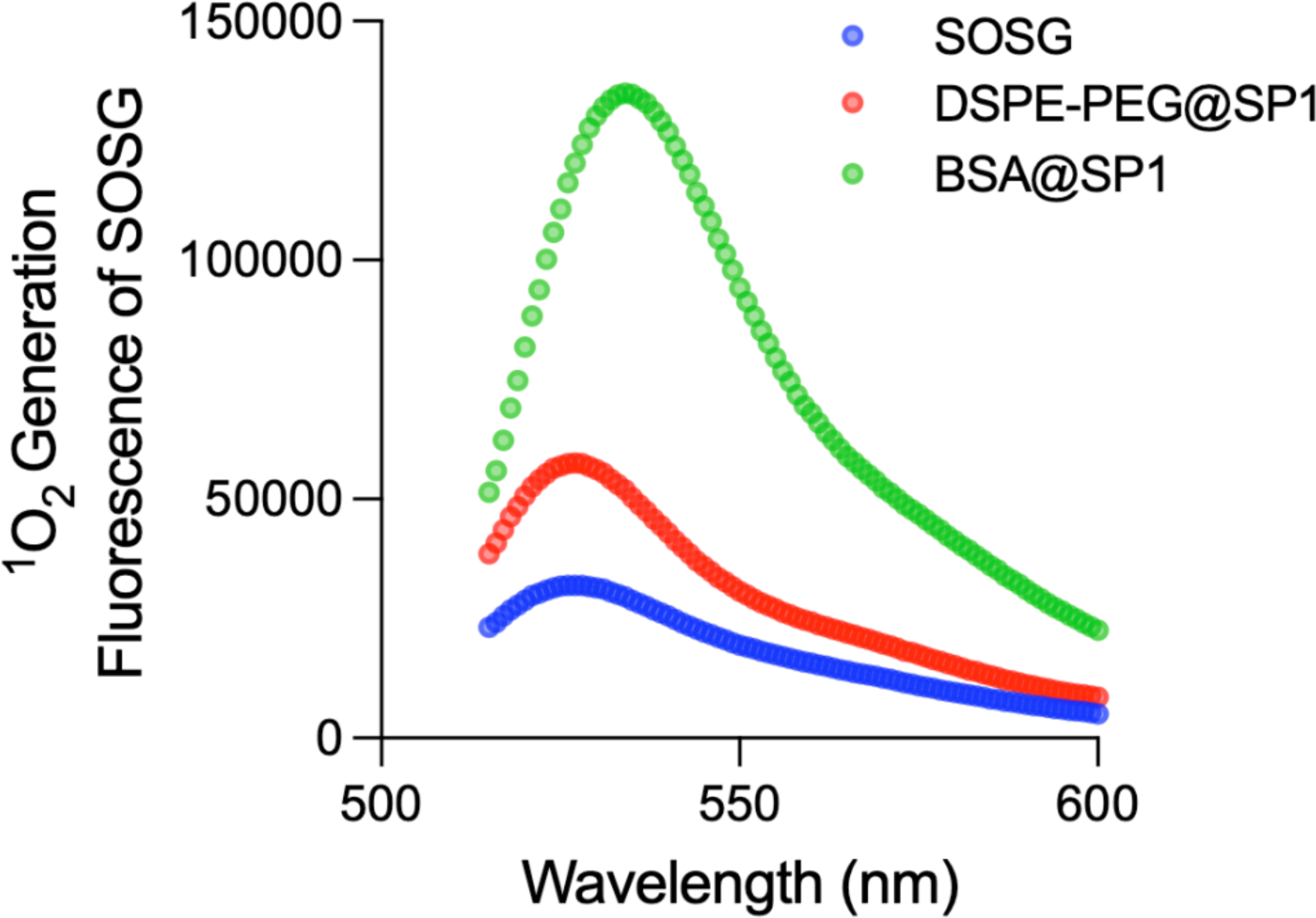
SOSG fluorescent spectra for samples treated either alone or with DSPE-PEG@SP1 or BSA@SP1 under white light irradiation for 60 seconds.

**Figure S18.**
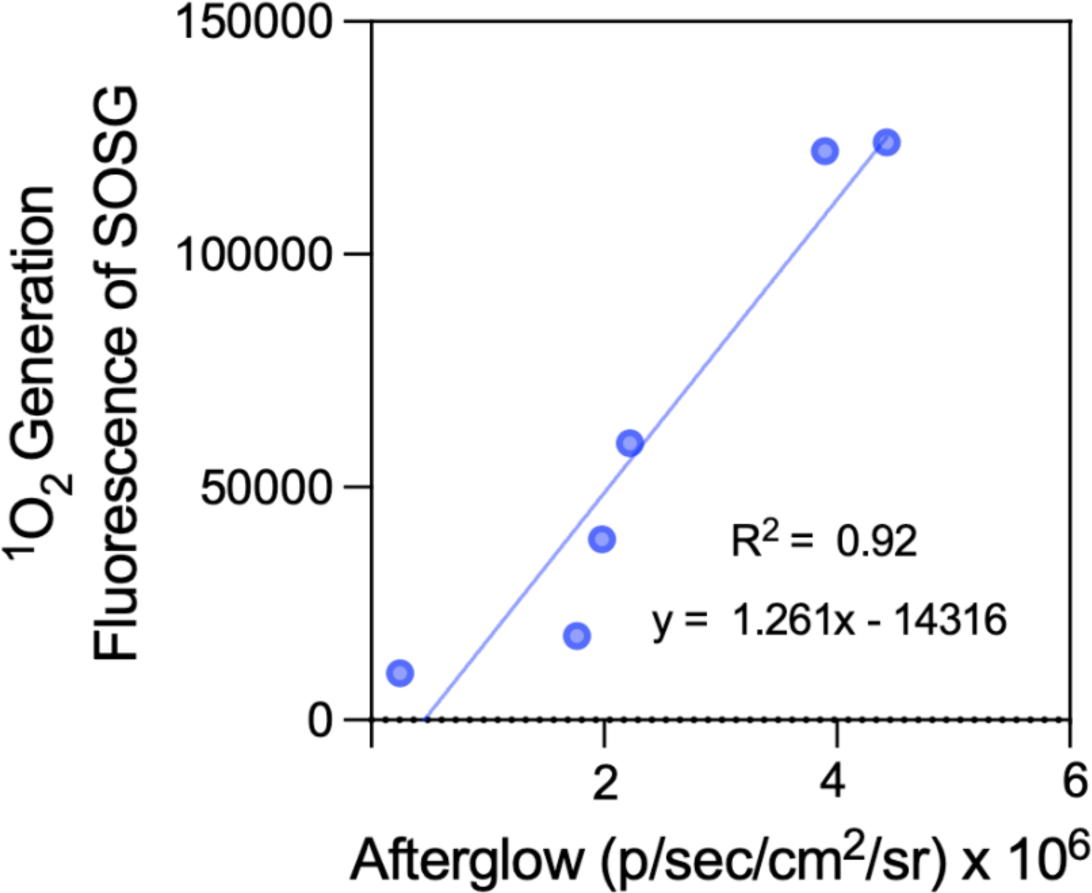
Correlation between afterglow intensity and ^1^O_2_ yield of BSA@SP1 calculated from IVIS images and SOSG detection.

**Figure S19.**
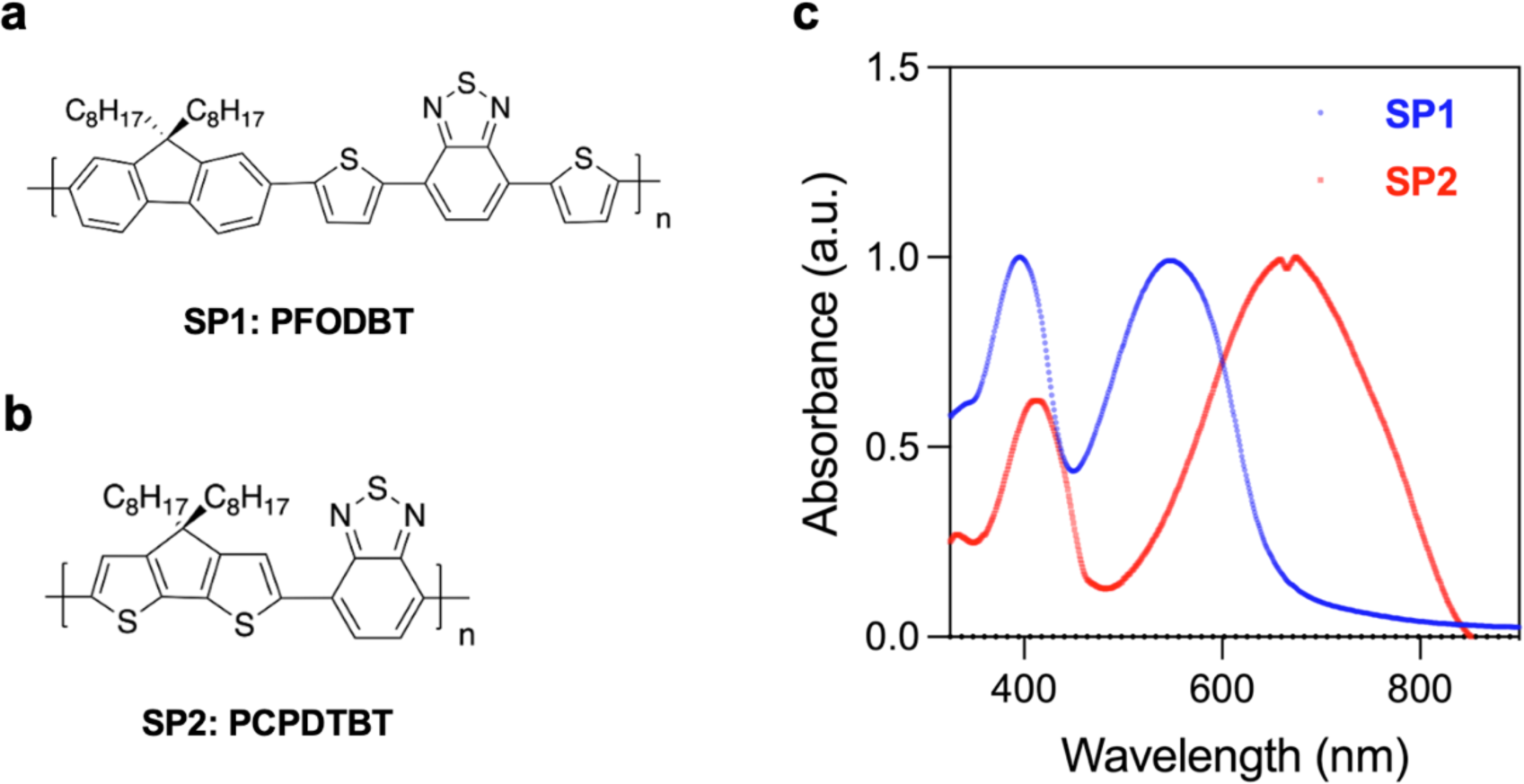
(a) Chemical structures of SP1 and SP2 and (b) their corresponding UV-VIS-NIR absorbance spectra.

**Figure S20.**
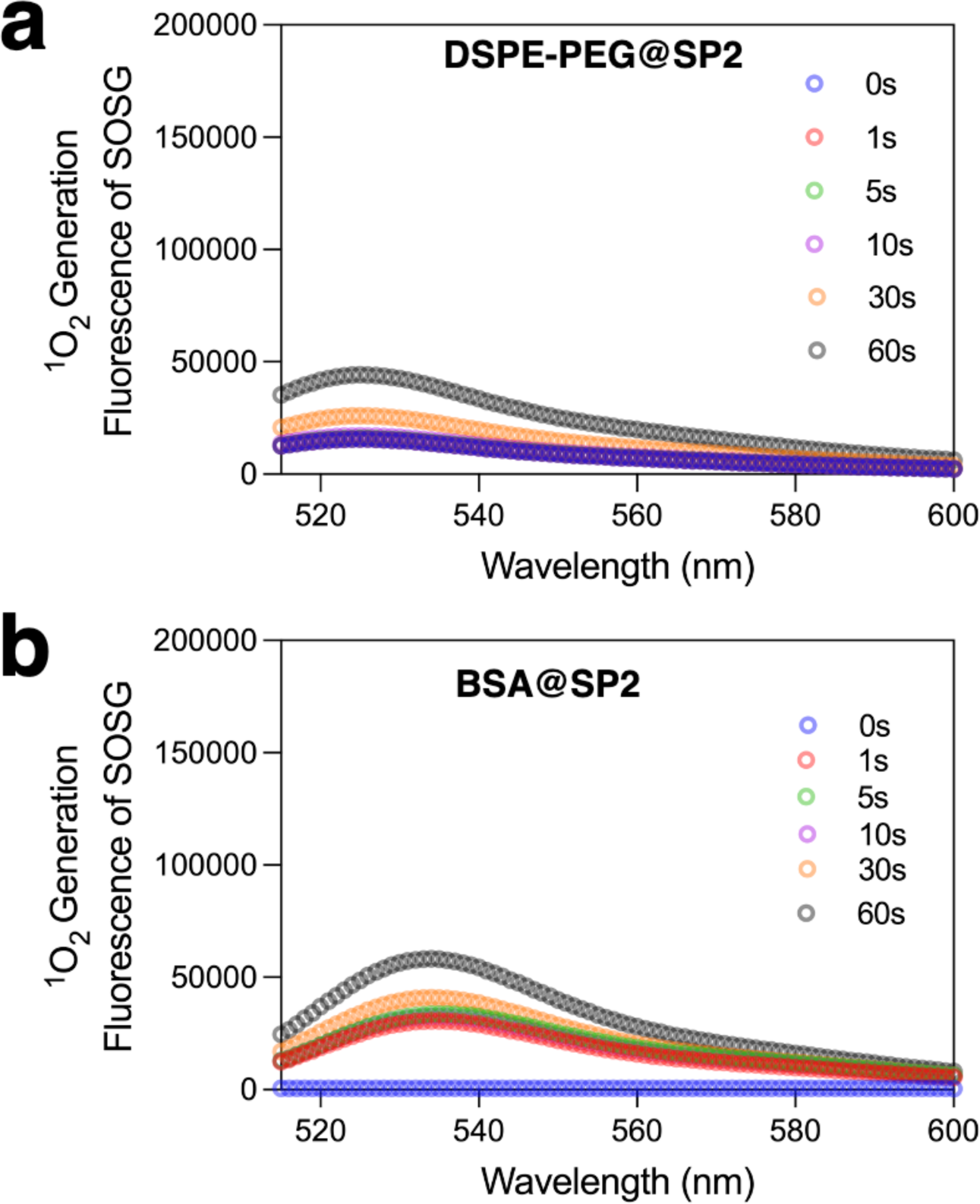
SOSG fluorescent spectra for (a) DSPE-PEG@SP2 and (b) BSA@SP2 correlated with their ^1^O_2_ generation treated with irradiations for different time durations.

**Figure S21.**
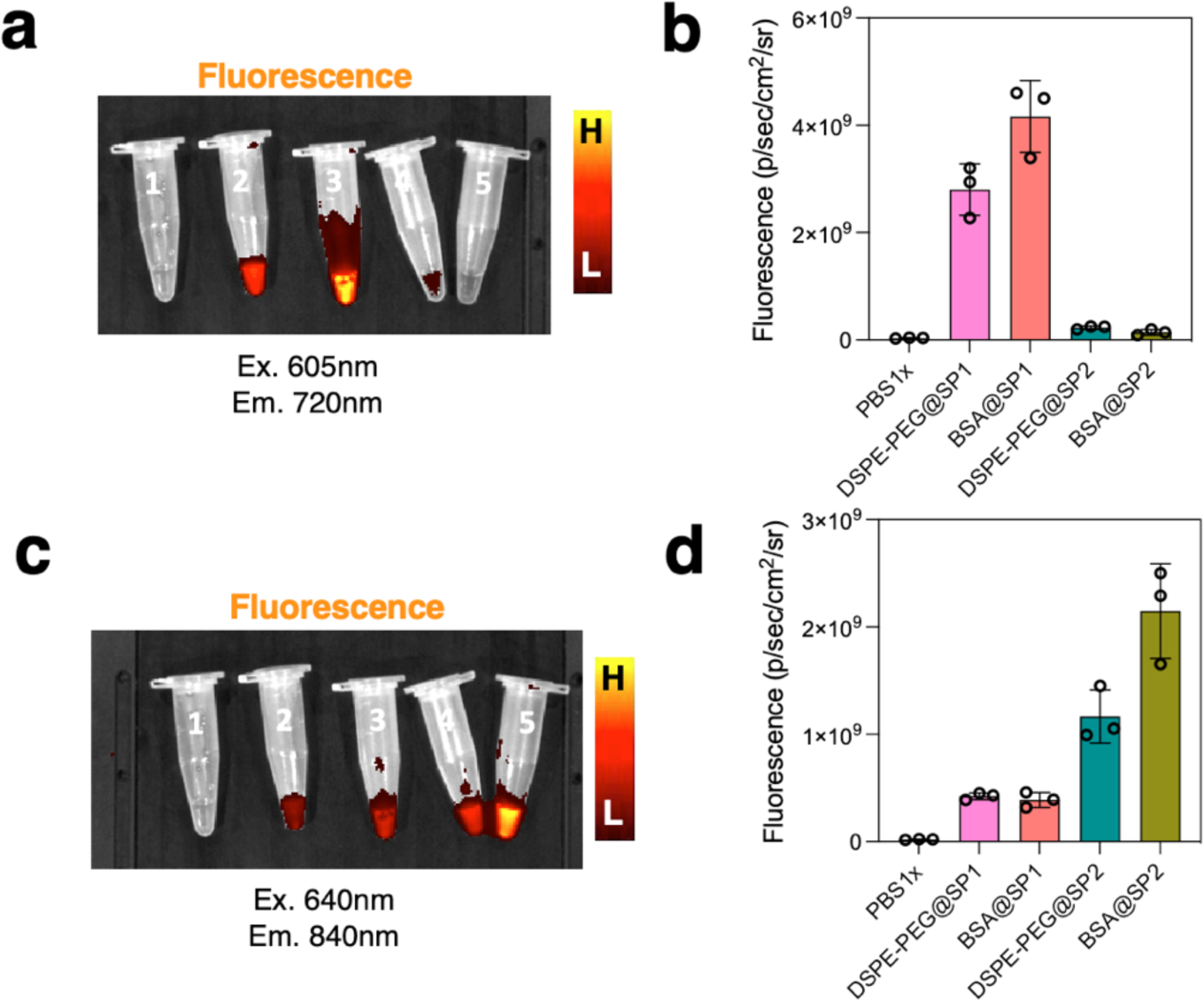
Fluorescence images obtained under (a) 605nm excitation, 720nm emission, (c) 640nm excitation, 840nm emission, for the following samples: (1) PBS1x, (2) DSPE-PEG@SP1, (3) BSA@SP1, (4) DSPE-PEG@SP2, (5) BSA@SP2. The corresponding fluorescence emission intensities were quantified in (b) and (d).

**Figure S22.**
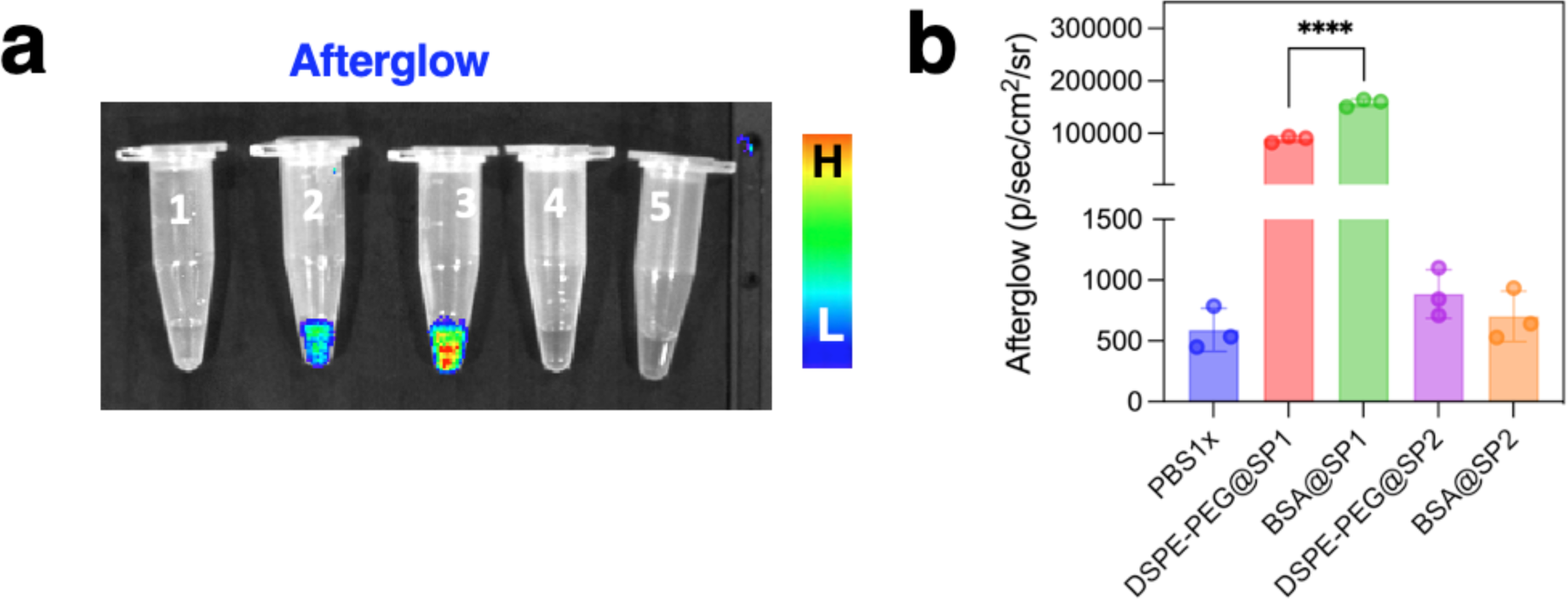
(a) Afterglow images were obtained under 1-minute white light irradiation and acquisition under an open filter in an IVIS imager for the following samples: (1) PBS1x, (2) DSPE-PEG@SP1, (3) BSA@SP1, (4) DSPE-PEG@SP2, (5) BSA@SP2. The corresponding afterglow emission intensities were quantified (b).

**Figure S23.**
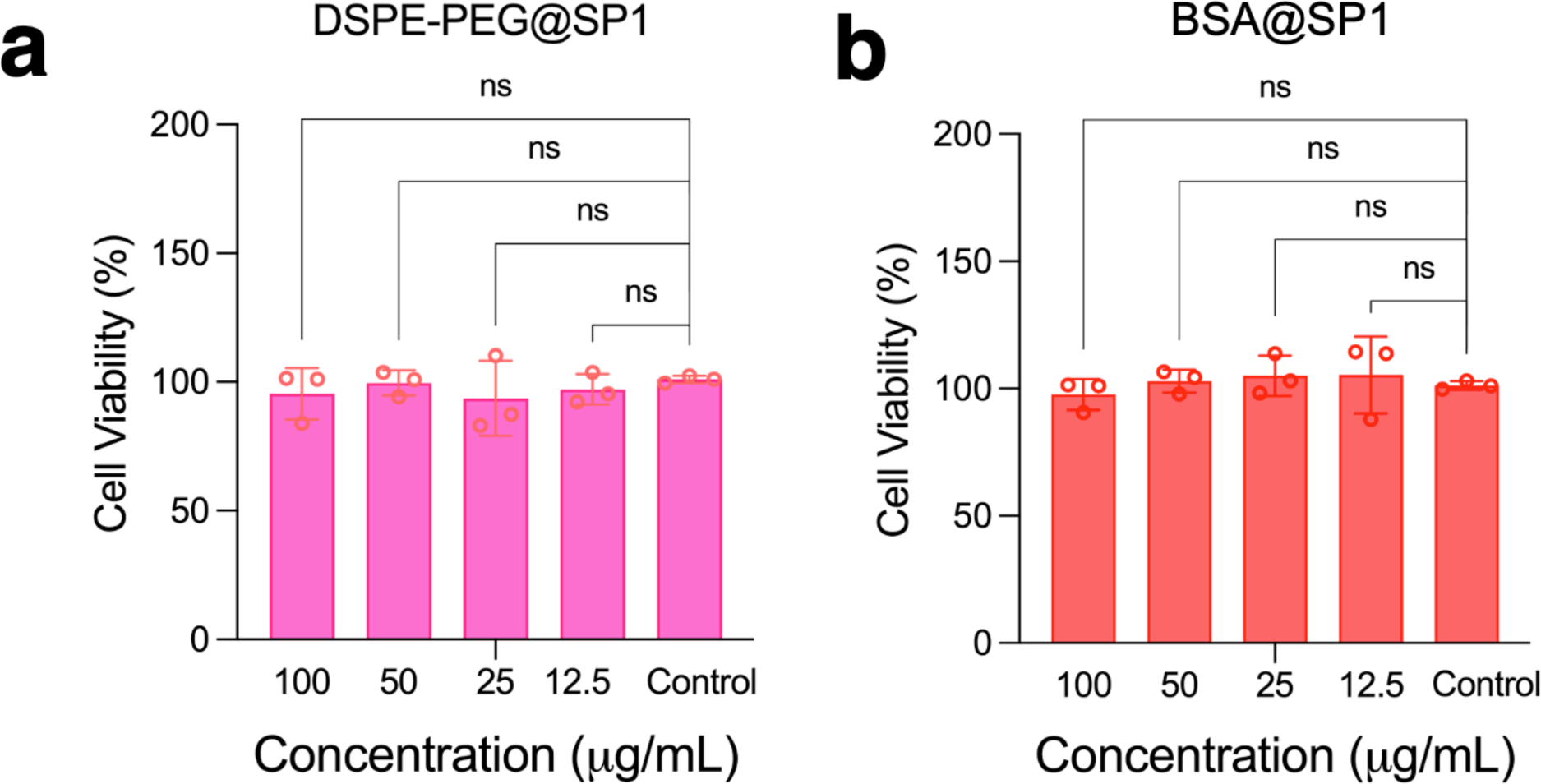
MTT cytotoxicity measurements of BSA@SP1 and DSPE-PEG@SP1 in 4T1 cells.

**Figure S24.**
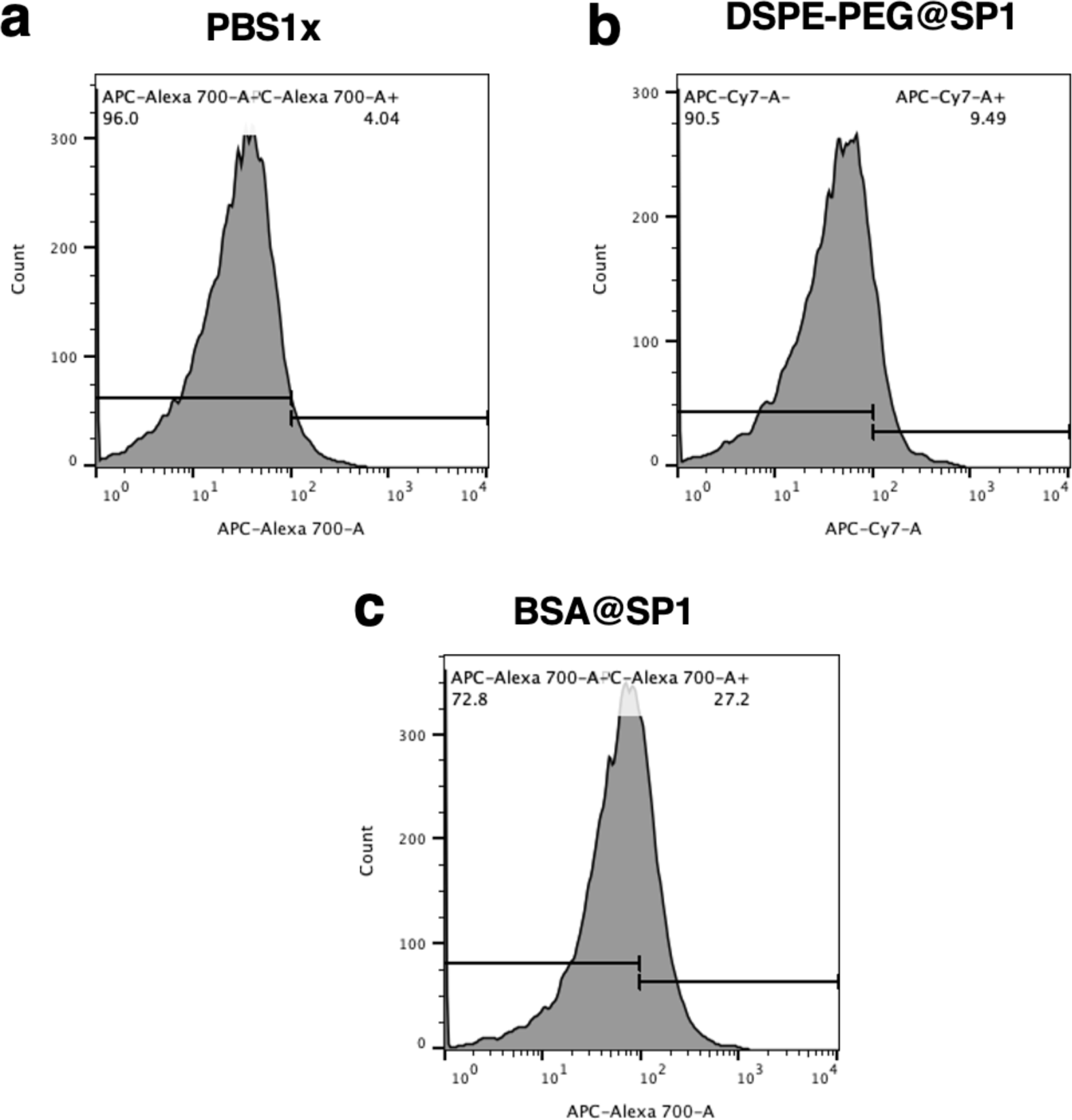
Cellular internalization analysis using representative flow cytometry plots for BSA@SP1 and DSPE-PEG@SP1 incubated in 4T1 cells for 8h.

**Figure S25.**
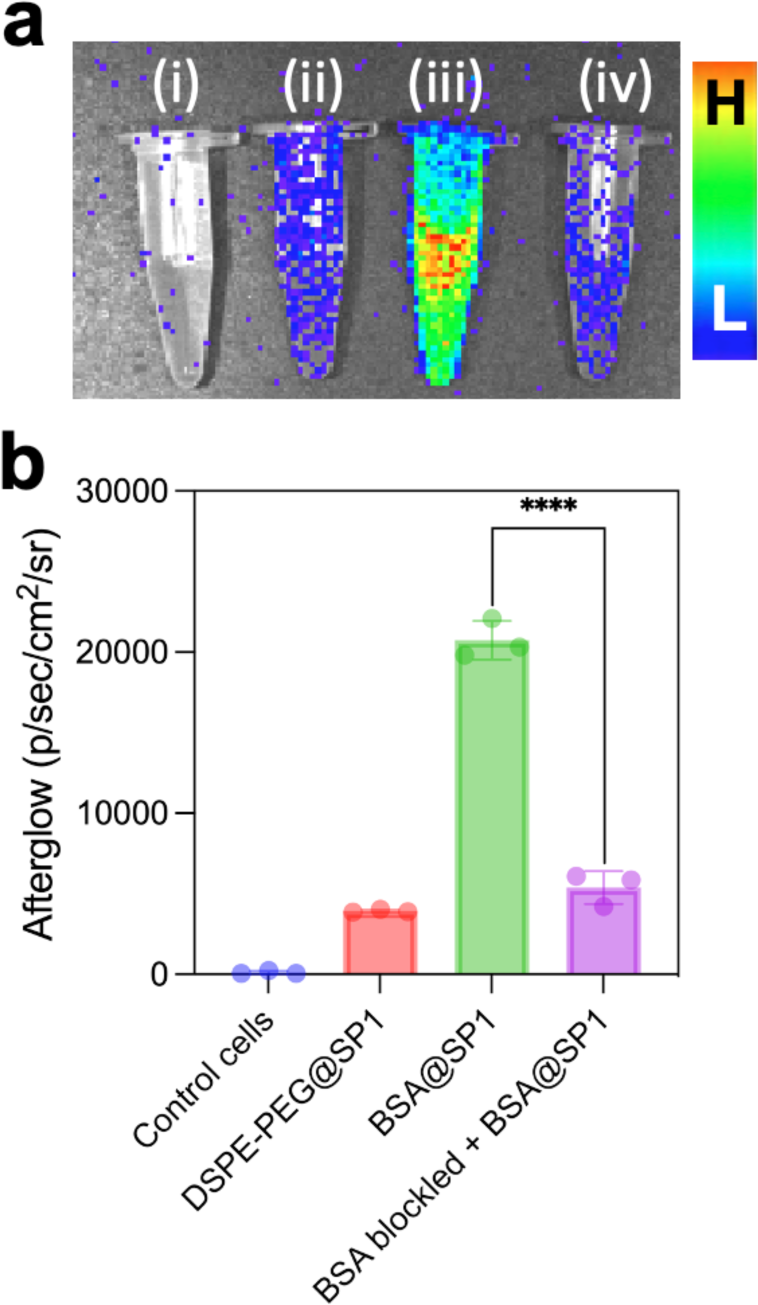
(a) Afterglow images were acquired in bioluminescence mode using an open filter for: (i) Control 4T1 cells, (ii) 4T1 cells incubated with DSPE-PEG@SP1, (iii) 4T1 cells incubated with BSA@SP1, and (iv) 4T1 cells pre-treated with 100 µM BSA solution, followed by BSA@SP1 treatment. All nanoparticle treatments were done at a concentration of 100 µg/mL for 4 hours. (b) Corresponding afterglow luminescence intensities for each condition.

**Figure S26.**
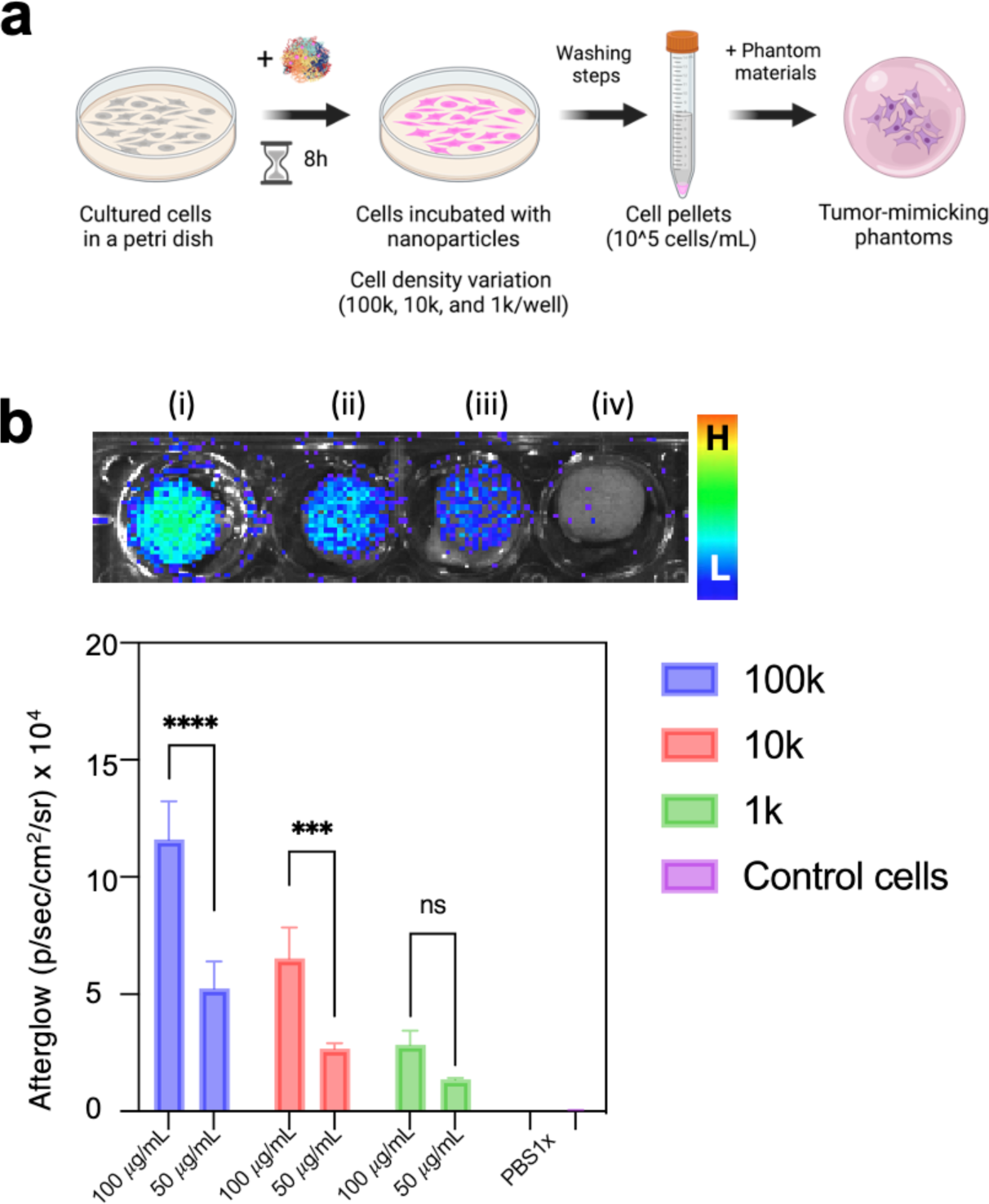
(a) A schematic representation illustrates the preparation of tumor cells-mimicking phantoms. These phantoms consist of BSA@SP nanoparticles internalized in 4T1 tumor cells, which are mixed with phantom solutions at varying cell densities of approximately 100,000, 10,000, and 1,000 cells, respectively. (b) Corresponding afterglow luminescence images were acquired using an open filter for the following conditions: (i) 100,000 cell density, (ii) 10,000 cell density, (iii) 1,000 cell density, and (iv) control cells pre-treated with BSA@SP1 at 100 µg/mL for 8 hours. The afterglow luminescence intensities were then compared across the different cell densities for two treatment concentrations (100 µg/mL and 50 µg/mL).

**Figure S27.**
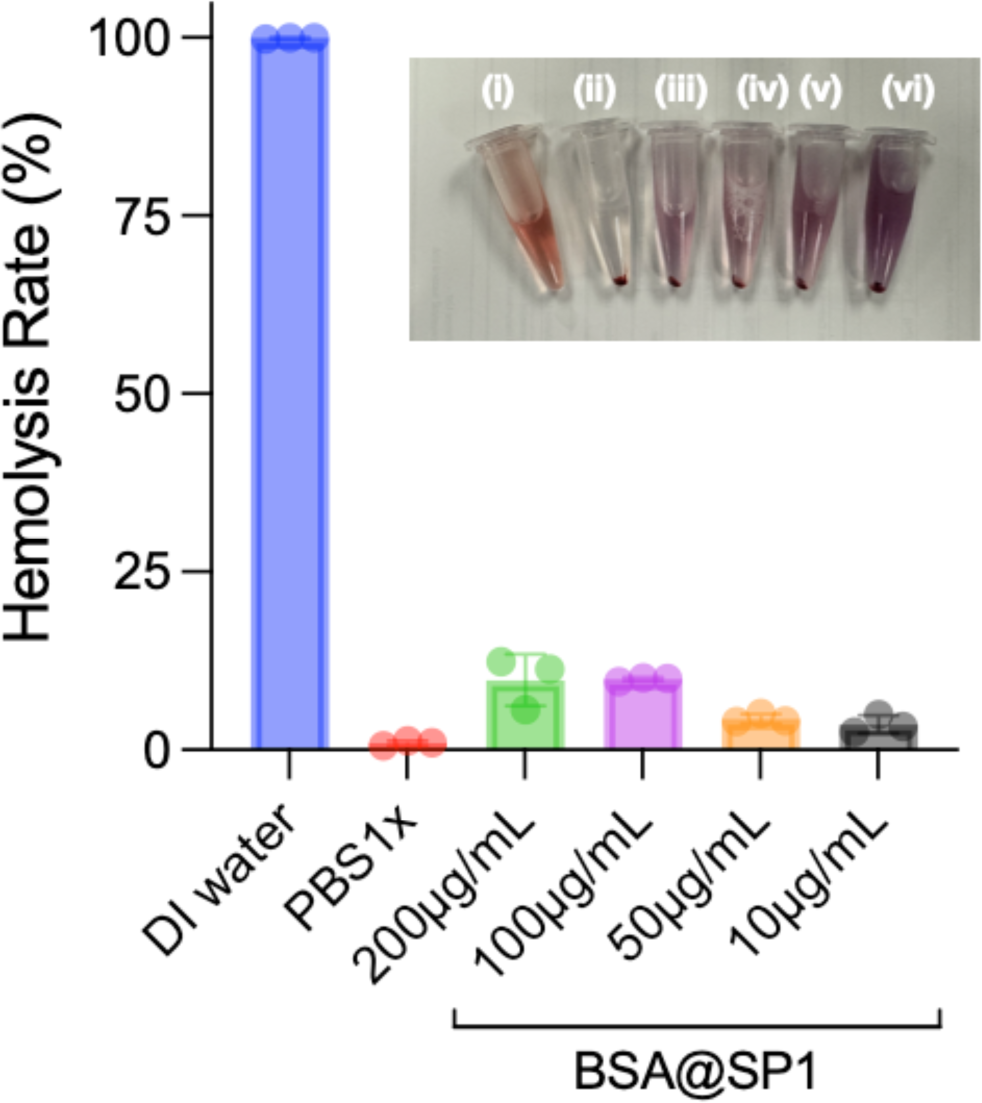
Hemocompatibility measurement of BSA@SP1 nanoparticles at different concentrations (10μg/mL, 50μg/mL, 100μg/mL, and 200μg/mL). PBS1x and DI water were taken as negative and positive controls, respectively.

